# Spatially and non-spatially tuned hippocampal neurons are linear perceptual and nonlinear memory encoders

**DOI:** 10.1101/2025.06.23.661173

**Authors:** Maxime Daigle, Kaicheng Yan, Benjamin Corrigan, Roberto Gulli, Julio Martinez-Trujillo, Pouya Bashivan

## Abstract

The hippocampus has long been regarded as a neural map of physical space, with its neurons categorized as spatially or non-spatially tuned according to their response selectivity. However, growing evidence suggests that this dichotomy oversimplifies the complex roles hippocampal neurons play in integrating spatial and non-spatial information. Through computational modeling and in-vivo electrophysiology in macaques, we show that neurons classified as spatially tuned primarily encode linear combinations of immediate behaviorally relevant factors, while those labeled as non-spatially tuned rely on nonlinear mechanisms to integrate temporally distant experiences. Furthermore, our findings reveal a temporal gradient in the primate CA3 region, where spatial selectivity diminishes as neurons encode increasingly distant past events. Finally, using artificial neural networks, we demonstrate that nonlinear recurrent connections are crucial for capturing the response dynamics of non-spatially tuned neurons, particularly those encoding memory-related information. These findings challenge the traditional dichotomy of spatial versus non-spatial representations and instead suggest a continuum of linear and nonlinear computations that underpin hippocampal function. This framework provides new insights into how the hippocampus bridges perception and memory, informing on its role in episodic memory, spatial navigation, and associative learning.

## 1 Introduction

The hippocampus is central to a diverse range of cognitive functions, including episodic memory formation (Scoville and Milner, 1957) and spatial navigation (O’Keefe, 1978). A key component of the hippocampus is its closed-loop neural structure that integrates multimodal inputs from diverse brain areas and projects its outputs to regions critical for memory, navigation, and decision-making (Buzśaki, 1996; Kesner and Rolls, 2015; Van Hoesen, 1982; Eichenbaum, 2017b). Traditionally, hippocampal function has been explored through two distinct lenses: its role in encoding and retrieving episodic memories and its function as a cognitive map for spatial navigation (Eichenbaum, 2004; O’Keefe, 1978). Early studies, particularly those exploring neural activity in the hippocampal formation during free navigation, identified spatially selective neurons such as place cells (O’Keefe and Dostrovsky, 1971) and grid cells (Hafting et al., 2005), which fire when a subject is at specific positions in space. These discoveries established the hippocampus as a “neural map” of the environment, a perspective that has profoundly influenced research on its function (Fig 1 H1, left).

**Figure 1:**
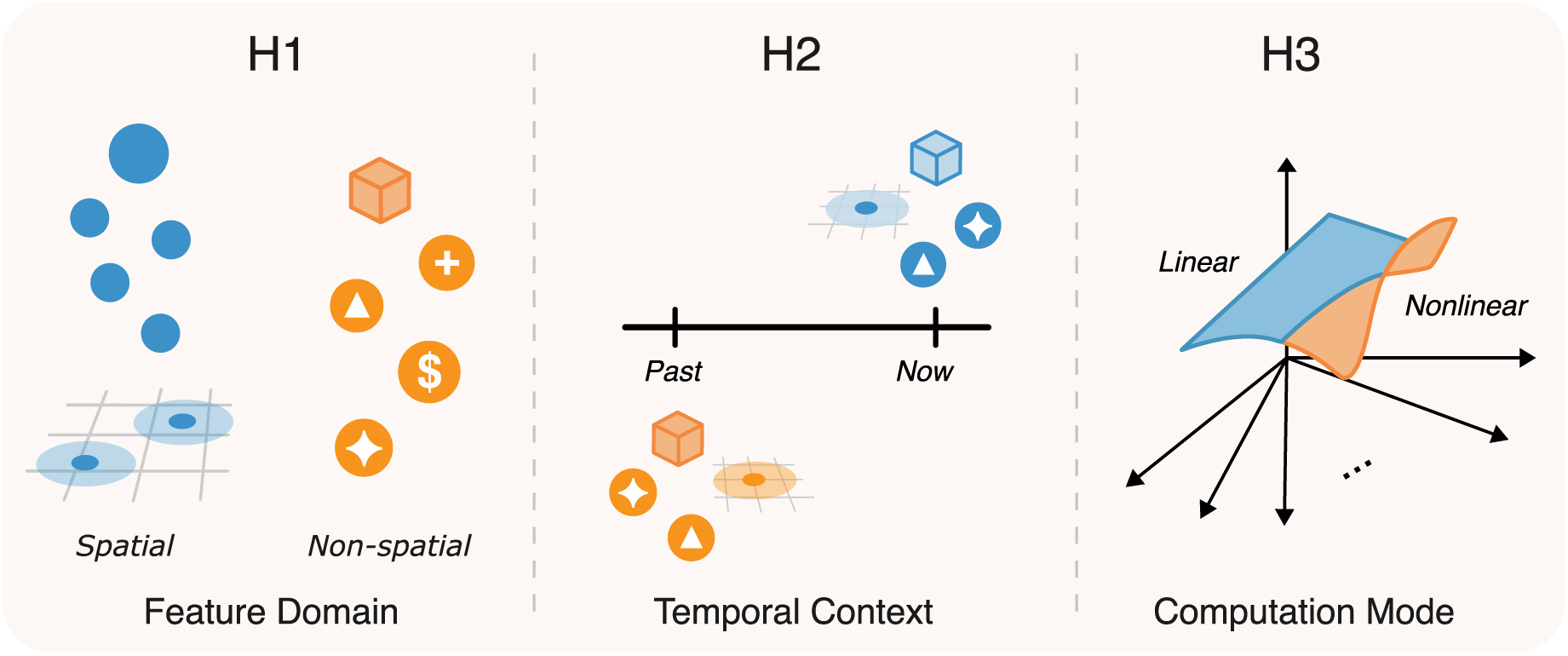
Hypothesized dimensions shaping hippocampal neural representation. While hippocampal investigations often focus on neuronal tuning to spatial or non-spatial features (H1 Feature Domain), a comprehensive understanding likely requires considering additional dimensions. Given the hippocampus’s established role in memory, the temporal dimension is hypothesized to be a critical factor shaping its neural representation (H2 Temporal Context). Concurrently, the nature of the underlying computational mechanisms, which may span linear to nonlinear operations, forms another key dimension (H3 Computation Mode). A more unified understanding of the hippocampus may thus depend on investigating how these different dimensions operate in concert.

More recent studies have expanded this view, showing that hippocampal neurons encode spatial and temporal dimensions of experience including space but also task-relevant features such as rewards (Gauthier and Tank, 2018), specific objects (Lin et al., 2007), and view-specific information (Rolls, 1999; Corrigan et al., 2023; Piza et al., 2024). These findings have fueled growing interest in characterizing both spatially and non-spatially tuned neurons (Eichenbaum et al., 1999; Zutshi et al., 2025) (Fig 1 H1, right), highlighting the diversity of representations found in the hippocampus.

This diversity of feature encoding naturally extends to the temporal domain, a critical dimension given the hippocampus’s well-established role in memory (Scoville and Milner, 1957). How does the hippocampus arbitrate between representing the immediate, unfolding environment and recalling experiences from the past (Eichenbaum, 2017a)? One possibility is that neurons specialize either in encoding current environmental features or in retrieving past experiences (Fig 1 H2, top vs. bottom). Alternatively, individual neurons may integrate inputs across temporal windows, giving rise to mixed-history representations.

Recent evidence shows that hippocampal neurons frequently encode a mixture of spatial and nonspatial features, such as item-place conjunctions (Komorowski et al., 2009), or integrate accumulated evidence with spatial position (Nieh et al., 2021). Spatial information, rather than being localized to specific neurons, is distributed across the hippocampal population. Moreover, single-neuron tuning only weakly predicts a neuron’s contribution to spatial encoding (Stefanini et al., 2020; Meshulam et al., 2017), raising questions about the functional roles of these tunings. In particular, they highlight our incomplete knowledge of the computational mechanisms underlying spatially and non-spatially selective neurons and how these mechanisms support the integration of spatial, temporal, and task-related information.

Recent computational modeling studies suggest that spatial selectivity in neurons can emerge as a byproduct of performing a range of ethologically relevant tasks, such as predicting future states or abstract structural relationships (Stachenfeld et al., 2017; Gornet and Thomson, 2024; Levenstein et al., 2024; Whittington et al., 2020), compressing sensory input (Benna and Fusi, 2021), and object recognition (Luo et al., 2024). A common principle uniting these models is that the observed tuning of a unit is a reflection of the underlying computational strategy it employs. While some representation might be adequately captured by linear combinations of variables, nonlinear operations enable the formation of complex neural representations (Rigotti et al., 2013). Characterizing the nature of these computational mechanisms and how they give rise to such flexible representations remains a key open question (Fig 1 H3).

To further probe these mechanisms, we combined computational modeling and in vivo electrophysiology in macaques to investigate the computational differences between spatially and non-spatially selective neurons. Using linear and nonlinear models, including reward-optimized artificial neural networks, we explored how hippocampal neurons encode information across spatial and temporal scales. Echoing prior findings (Stefanini et al., 2020; Meshulam et al., 2017), our results also demonstrate that spatial and non-spatial tuning do not imply exclusive encoding of spatial or non-spatial information by a neuron. Instead, they reveal that spatial tuning reflects a neuron’s tendency to encode linear combinations of task-relevant features, whereas non-spatially tuned neurons primarily encode nonlinear combinations of features. Moreover, we observed a parallel relationship between the linearity of encoding and mnemonic representations: neurons encoding currently perceived information are more readily explained by linear combinations of features, whereas those encoding past information rely more heavily on nonlinear encoding strategies. Notably, neurons become progressively less likely to exhibit spatial tuning as they encode information from further in the past.

These results suggest that memory neurons rely on nonlinear computation to integrate information across time and space. As the temporal distance from an event increases, spatial tuning becomes less prominent, and neuronal responses tend to reflect more abstract, nonlinear combinations of past inputs. This gives rise to a continuum of neuronal profiles, with spatially tuned neurons that linearly encode currently perceived information at one end, and non-spatially tuned neurons that nonlinearly encode past information at the other. This gradient suggests that the functional diversity in the hippocampus is shaped not by a spatial–temporal dichotomy, but by differences in the underlying computational mechanisms—ranging from linear to nonlinear integration—enabled by its recurrent circuitry. This graded shift in coding strategy is consistent with the hippocampus’s recurrent circuitry and its role in linking the present with events that are separated in time and space (Howard and Eichenbaum, 2015). This spectrum supports the formation of structured representations that generalize across experiences, providing a foundation for encoding complex relational memories that unfold over time.

## 2 Results

### 2.1 Spatially-tuned hippocampal neurons are largely mixed-selective

Prior works on mixed selectivity demonstrated that hippocampal place fields are modulated by factors such as view (Rolls, 1999; Corrigan et al., 2023; Piza et al., 2024), reward locations (Rolls and Xiang, 2005), and context (Anderson and Jeffery, 2003). With learning, some neurons develop a firing map that integrates goal-centered and task-related information (Baraduc et al., 2019). Additionally, the cognitive map of space does not generalize across tasks, even within the same environments, indicating that the representation of space is not invariant to contexts and suggesting that spatial representation may depend on the encoding of task-related features (Gulli et al., 2020). To investigate the computational role of spatial tuning in the primate hippocampus, we analysed 175 single-neuron recordings predominantly from the macaque CA3 sub-region of the hippocampus (Gulli et al., 2020). Macaque monkeys were seated in front of a computer monitor and used a joystick to complete an associative memory task that required navigating through a virtual maze (Fig 2a).

**Figure 2:**
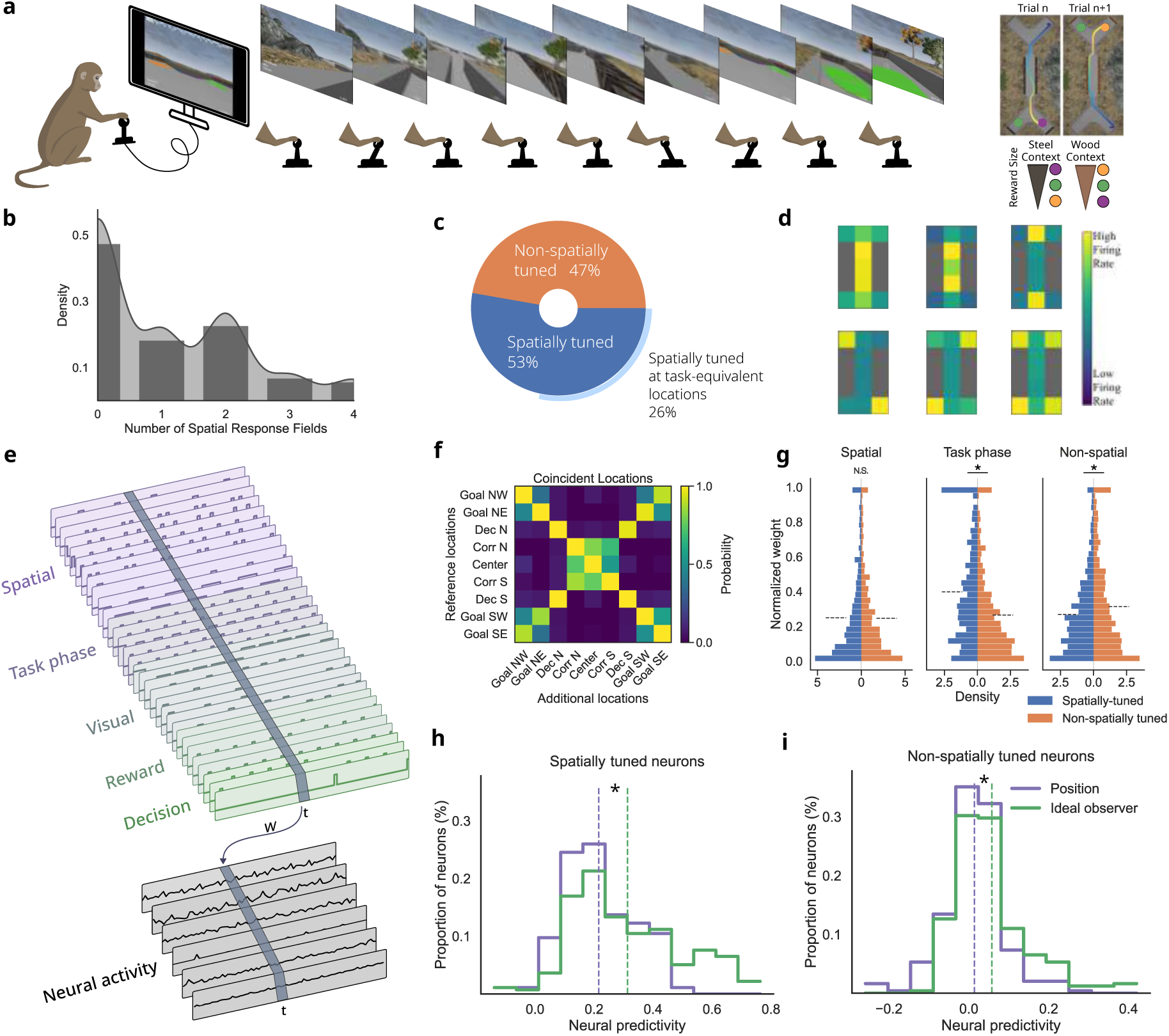
Encoding of spatial and non-spatial features. (a) Schematic representation of the navigation-dependent associative memory task performed by macaque monkeys in a virtual maze. (b) Distribution of spatial response fields per neuron. (c) Proportion of neurons exhibiting spatial tuning. Non-spatially tuned neurons do not exhibit any spatial response field. Spatially tuned neurons exhibit at least one. Spatially tuned at task-equivalent locations is the subset of spatially tuned neurons with spatial response field distributed in patterns demonstrated in c. (d) Examples of spatial map for six STNs with multiple response fields. (e) Schematic of the ideal observer model used to predict hippocampal neuron firing rates based on task-relevant features. (f) Coincident locations of spatial response fields for neurons with multiple response fields, demonstrating symmetrical tuning properties. (g) Distribution of fitted weights from the ideal observer model across neuron types. Histograms show normalized weights for spatial (left), task phase (center), and non-spatial (right) features, comparing spatially tuned versus non-spatially tuned neurons. Dashed lines represent the mean weight. (Wilcoxon rank-sum test: non-significant difference between spatial weights in spatially and non-spatially tuned neurons *P* = 0.14; greater weights on task phase features in spatially tuned neurons *P <* 10*^−^*^34^; greater weights on non-spatial features in non-spatially tuned neurons*P <* 10*^−^*^6^). (h-i) Histogram of neural predictivity scores comparing two model conditions for spatially tuned and non-spatially tuned neurons. Mean neural predictivity scores indicated by vertical dashed lines. (Wilcoxon rank-sum test: STNs *P <* 10*^−^*^9^, nSTNs *P <* 10*^−^*^4^). All regressions are repeated across three random seeds. Ideal observer contains all task-relevant features, while the position model only contains position features. N, north; S, south; W, west; E, east. Mean *±* standard deviation over three random seeds

The distribution of spatial response fields across neurons was bimodal: while most neurons exhibited no spatial response fields, many spatially tuned neurons (STNs) had two spatial response fields (Fig 2b). Neurons with multiple spatial response fields tended to exhibit elevated firing rates at multiple locations that hold functional equivalence within the task structure, manifesting as symmetrical tunings (Fig 2d,f). Even though 53 *±* 1% of hippocampal neurons exhibited at least one spatial response field, 49 *±* 2% of spatially-tuned neurons have their fields distributed across task-equivalent locations (Fig 2c; Fig S1). This suggests that while those neurons show spatial tuning, their encoding is not purely spatial, but rather reflects a complex integration of multiple task-relevant features.

The hippocampus serves as a sensory hub that receives a variety of highly processed information, encompassing spatial, temporal, visual, olfactory, and auditory stimuli (Eichenbaum, 2017a; Save et al., 2000; Itskov et al., 2012) which puts the hippocampus in a unique position for representing information across modalities. To investigate whether spatial tuning property of neurons is indicative of whether a neuron encodes a single feature or a mixture of them, we utilized a linear ideal observer model to predict the rate of spiking activity in each recorded neuron from all task-relevant features including spatial position, direction, visual cues, decision, and reward (Fig 2e). To understand the relative importance of spatial and non-spatial features for each group of neurons, we quantified the contribution of each type of feature through its corresponding fitted weight. While non-spatially tuned neurons (nSTNs) relied significantly more on non-spatial features (Fig 2g greater weights on non-spatial features in non-spatially tuned neurons, Wilcoxon rank-sum test: *P <* 10*^−^*^7^), STNs exhibited strong encoding of task phases (Fig 2g task phases, Wilcoxon rank-sum test *P <* 10*^−^*^35^). Interestingly, weights on spatial features did not differ significantly between STNs and nSTNs (Fig 2g spatial, Wilcoxon rank-sum test: *P* = 0.14), which corroborate with both the strong encoding of task phases by STNs (which are partly correlated with spatial position) and previous findings indicating that nSTNs also contribute to position encoding in CA1 (Stefanini et al., 2020; Meshulam et al., 2017).

Moreover, the ideal observer model improved the prediction accuracy of both spatially and non-spatial tuned neurons responses, beyond those from the spatial position-only regression model, suggesting a mixed-selective coding scheme (Fig 2h-i). Interestingly, spatially-tuned neurons benefited more from linear combination of non-spatial features than non-spatially tuned neurons (Fig 2h-i, ideal observer better predicts neural responses than position model, Wilcoxon rank-sum test: STNs *P <* 10*^−^*^9^, nSTNs *P <* 10*^−^*^4^; Fig S2, the gap between both models was smaller for nSTNs compared to STNs, two-sample Kolmogorov–Smirnov test *P <* 10*^−^*^12^).

### 2.2 Spatially and non-spatially tuned neurons encode linear and nonlinear combinations of task-relevant features

Many hippocampal neurons exhibit context-dependent response patterns that may not be easily simulated with linear models, such as the ideal observer model (Anderson and Jeffery, 2003). A promising choice of nonlinear features are those that are useful for solving the given task. We reasoned that a minimally unbiased set can be automatically constructed by training an Artificial Neural Network (ANN) to perform the virtual navigation task to maximize its return (Fig 3a-b). We trained the Reward Optimized Artificial Recurrent Network (ROARN) to perform a virtual navigation task from egocentric sensory observations. The ANN architecture consisted of a convolutional network followed by a gated recurrent network. The virtual navigation task was simulated in a 2D environment, mirroring the environment and the task used in the animal experiment (Fig 3b). Consistent with the paradigm used for training the animal subjects, the ANN model received tiered rewards for navigating to context-dependent goals.

**Figure 3:**
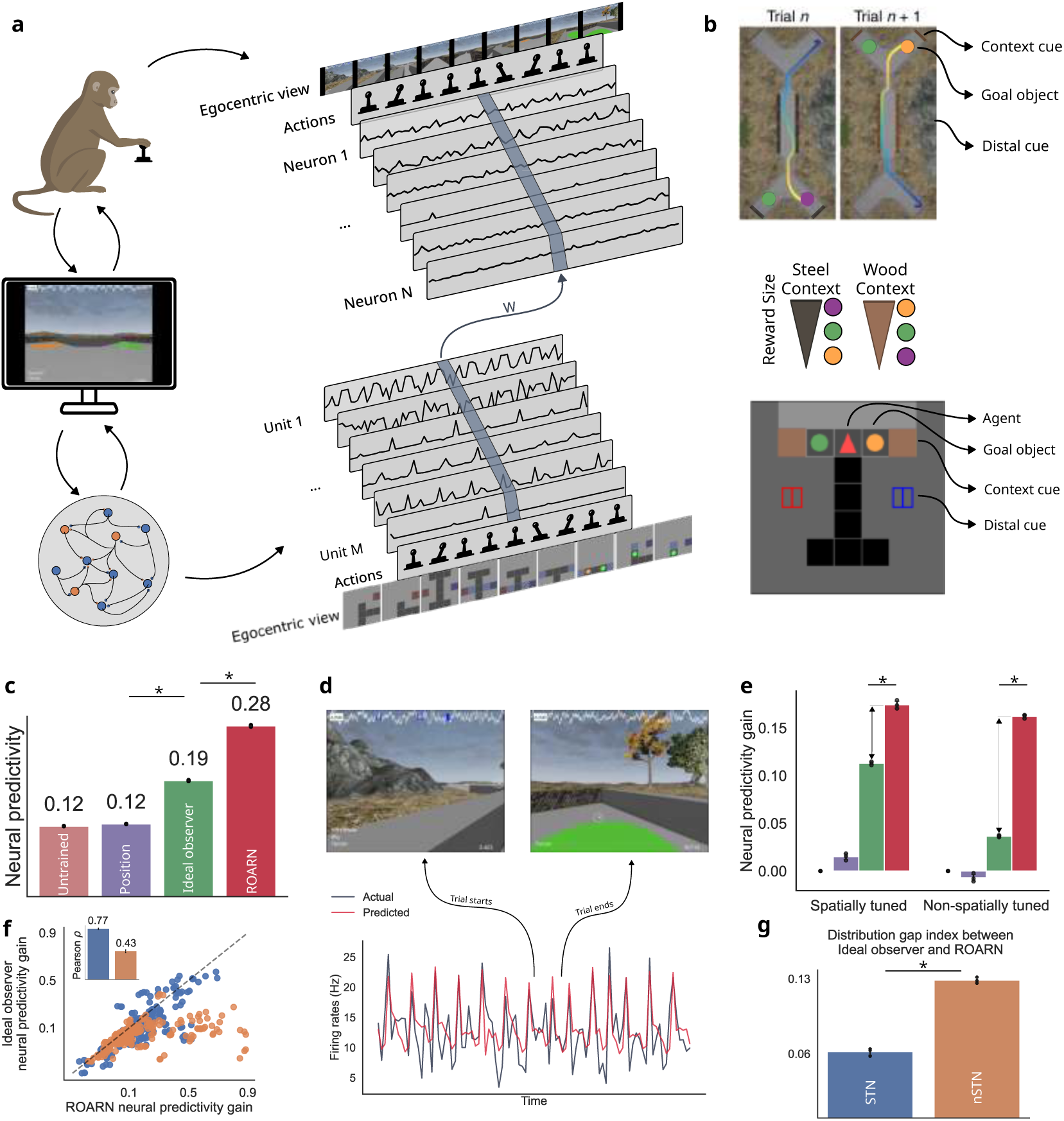
Predicting neural responses using computational models. (a) Schematic representation of the virtual navigation task performed by macaque monkeys and artificial agents, and the prediction of neural responses. Task performed by monkey is reproduced in an environment allowing ANN models to perform the same associative memory task. While both the monkey and artificial agent perform the same sequence of trials, neuronal responses are aligned with the ANN activations. A linear mapping is learned to transform the responses from the artificial units space to the neuronal space and allows to measure the similarity between both. (b) Top-down view of the maze for the associative memory task. Top – Monkey’s environment, Bottom – Agent’s environment, Middle – Example of reward schema for a session. The goal object providing the best reward in a context is the worst goal in the other context. The goal colors and their reward value are randomly changed at each session and the animals / agents need to learn the new association. (c) Neural predictivity of each models across all neurons, Wilcoxon rank-sum test: Position’s neural predictivity is less than the Ideal observer model *P <* 10*^−^*^6^ and Ideal observer’s neural predictivity is less than ROARN *P <* 10*^−^*^11^. (d) Example of hippocampal neuron responses and their corresponding predictions from the ROARN model. (e) Comparison of neural predictivity gain across models per subset of neurons, Wilcoxon rank-sum test: spatially *P <* 10*^−^*^9^ and non-spatially tuned neurons *P <* 10*^−^*^10^. (f) Scatter plot of neural predictivity gain of ROARN and ideal observer. (g) The gap in neural predictivity gain between ROARN and ideal observer increases for nSTNs (Wilcoxon rank-sum test: *P* = 0.02). All regressions were repeated across three random seeds. Mean *±* standard deviation over three random seeds

Following training, we constructed a predictive model of CA3 neural responses by fitting a linear mapping function to predict neural activity at each site using the activity of ROARN’s artificial units as predictors (Fig 3d; Fig S3). Despite relying on raw sensory inputs rather than abstract task relevant features like spatial position and object classes, this model outperformed both linear models at predicting CA3 neural responses (Fig 3c, Wilcoxon rank-sum test: Position predicts neural responses less well than Ideal observer model *P <* 10*^−^*^6^ and Ideal observer’s neural predictivity is less than ROARN *P <* 10*^−^*^11^).

However, the relative improvement in prediction accuracy between ROARN and the ideal observer model varied across neurons. A significant portion of nSTNs were significantly better predicted by the ROARN model (Fig 3e nSTNs, Wilcoxon rank-sum test *P <* 10*^−^*^10^). In contrast, the neural predicitivity was more comparable between the ideal observer and ROARN models for STNs (Fig 3e STNs, Wilcoxon rank-sum test *P <* 10*^−^*^9^). The improvement in neural predictivity for nSTNs was approximately twice that of STNs (Fig 3f,g; Wilcoxon rank-sum test *P* = 0.02). As a result, the gap between ROARN and ideal observer widened considerably for nSTNs, suggesting a greater reliance on nonlinear computations in those neurons.

These findings suggest that the long-standing debate over the function of spatial and non-spatial tuning in hippocampal neurons may be better understood through the computational functions these neurons implement. Specifically, STNs predominantly encode information through linear combinations of spatial and non-spatial features, whereas nSTNs rely on nonlinear combinations. This indicates that the underlying representation in macaque CA3 is not inherently space-based but instead spans a spectrum of linear and nonlinear mixtures of spatial and non-spatial features. Consequently, neurons classified as spatially tuned may simply be those that employ linear functions of these features, making traditional spatial selectivity tests more sensitive to detecting variations in neural activity based on spatial position (Behrens et al., 2018).

### 2.3 Non-spatially tuned neurons rely on nonlinear recurrent computations

The ROARN and ideal observer models represent fundamentally different computational frameworks, with several key factors contributing to their differences. First, the models receive distinct inputs: ROARN processes egocentric visual information, while the ideal observer is provided with abstract, task-relevant features. Second, ROARN’s representations emerge through task optimization as it learns to perform the navigation-dependent associative memory task, whereas the ideal observer lacks this learning-driven process. Third, the models differ in their dimensionality. Fourth, ROARN incorporates nonlinear transformations, while the ideal observer is constrained to linear combinations. Finally, ROARN includes a recurrent layer, enabling dynamic information integration over time, whereas the ideal observer does not.

We hypothesized that the nonlinear recurrence in the ROARN model was critical for its enhanced similarity to hippocampal neural activity. To test this (Fig 4a), we designed a modified version of ROARN with linear recurrence (ROARN w/ LR), replacing the model’s nonlinear recurrent gating architecture with a linear recurrence while keeping all other components—including nonlinear transformations in other layers—unchanged. The modified ROARN maintained its ability to learn and perform the task at the same proficiency as the original model (Fig S4). However, this change abolished ROARN’s advantage in predicting nSTN responses (Fig 4c, d), reducing its predictive performance to the level of the ideal observer for STNs and nSTNs (Fig 4b,c; Wilcoxon rank-sum test: no significant differences for STNs and nSTNs; Fig S5). These findings underscore the essential role of nonlinear recurrence in enabling ROARN’s superior neural predictivity compared to the ideal observer.

**Figure 4:**
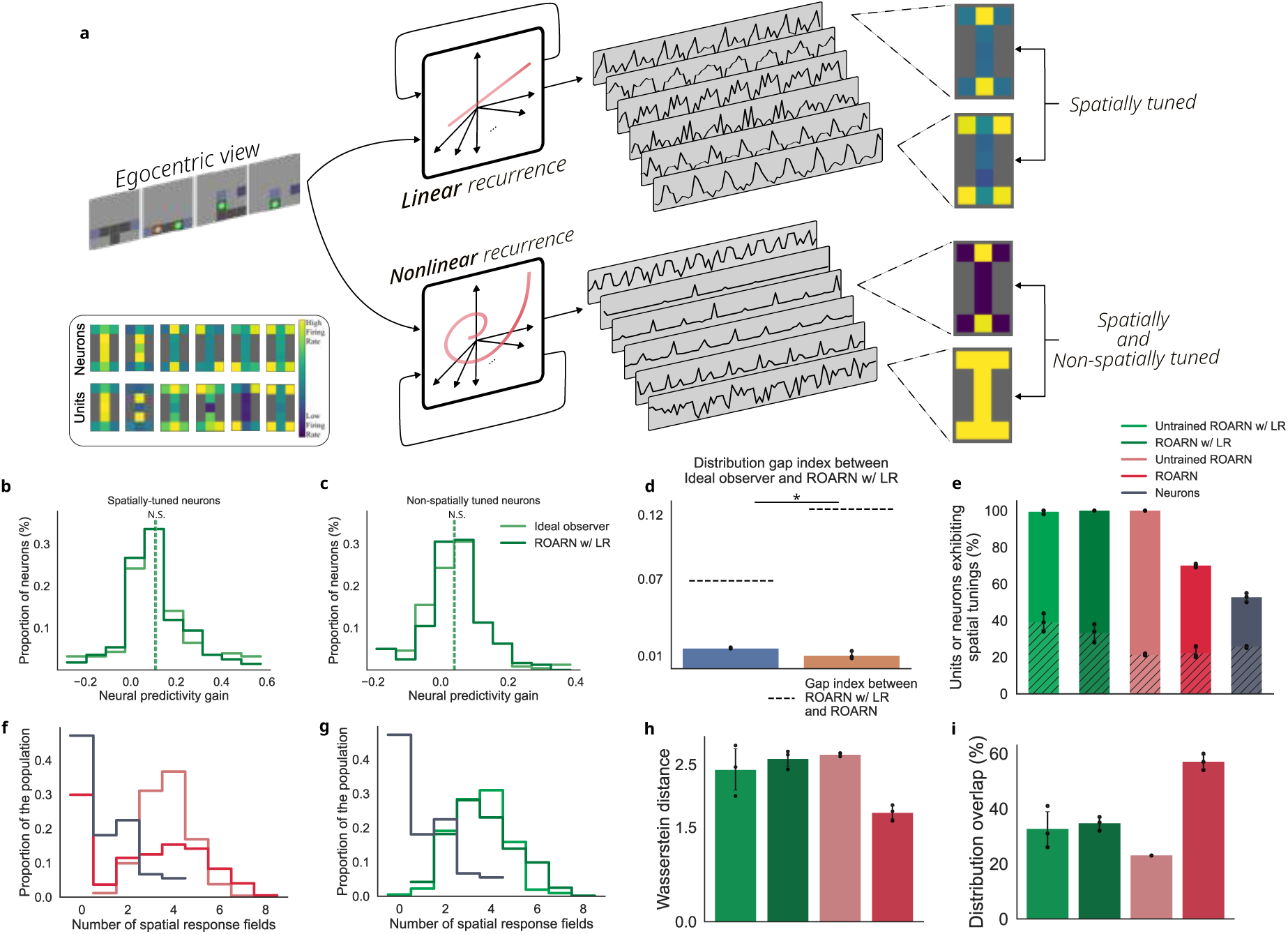
Non-spatially tuned neurons rely on nonlinear recurrent computations. (a) Schematic illustrating the relationship between spatial tuning and nonlinear recurrent encoding. Bottom left inset: example spatial response maps observed in spatially tuned neurons and artificial units from ROARN. (b-c) Histogram of neural predictivity gain comparing ROARN w/ LR and the ideal observer for spatially tuned (STNs) and non-spatially tuned neurons (nSTNs). Dashed lines indicate mean neural predictivity (Wilcoxon rank-sum test: *P* = 0.60 and *P* = 0.29 respectively). (d) Neural predictivity gain gap index between the ideal observer and ROARN w/ LR, showing a small gap for both STNs and nSTNs. Dashed lines indicate the predictivity gap between ROARN and ROARN w/ LR, which was approximately twice as large for nSTNs compared to STNs (Wilcoxon rank-sum test: *P* = 0.02). (e) Proportion of spatially tuned neurons and artificial units from ROARN and ROARN w/ LR. Hatched regions indicate the fraction of spatially tuned neurons/units with multiple response fields distributed across task-equivalent locations. (f) Distribution of spatial response fields in ROARN before and after training. A third of initially spatially tuned units lost their tuning after task optimization, becoming non-spatially tuned. (g) Spatial response field distribution in ROARN w/ LR before and after training, showing minimal change after learning the task. (h) Wasserstein distance between the distributions of spatial response fields in artificial units and neurons, where a lower distance indicates greater similarity to hippocampal spatial tuning distributions. (i) Overlap between the spatial response field distributions of artificial units and hippocampal neurons (f, g), with higher overlap indicating better alignment with neuronal recordings. All regressions were repeated across three random seeds. Mean *±* standard deviation over three random seeds

We further investigated whether spatial tuning properties emerged in artificial units of these ANNs in a manner analogous to those observed in primate hippocampal neurons (Fig 4a). Remarkably, we found that, before task optimization, all untrained units in both the ablated and original models exhibited spatial tuning, likely due to distinct input patterns across spatial locations, which generated unique activation profiles per location. However, task optimization had divergent effects depending on the presence of nonlinear recurrence. Intriguingly, training ROARN w/ LR did not substantially alter the distribution of spatial response fields (Fig 4e, green bars; Fig S6-S8). In contrast, the presence of nonlinear recurrence facilitated a transformation wherein some spatially tuned units evolved into non-spatially tuned units (Fig 4e, red bars; Fig S7). This resulted in a bimodal distribution of spatial response fields—one mode with no spatial tuning and another centered around four fields (Fig 4f). Without nonlinear recurrence, the training process failed to produce this second mode of non-spatially tuned units (Fig 4g). Consequently, nonlinear recurrence was necessary to generate a distribution more closely resembling that of the hippocampus (Fig 4h,i; Fig S9). This shift is consistent with findings in rodents, where spatial tunings are observed during initial exploration of an environment, but conjunctive coding gradually emerges with experience (Navratilova et al., 2012). Together, these results suggest that nonlinear recurrent connections are crucial for nSTNs, not only for the emergence of non-spatially tuned units, but also for accurately modeling the neural responses of nSTNs.

### 2.4 Nonlinearity of mnemonic CA3 neural code

This dichotomy between spatially and non-spatially tuned neurons, with the former favoring linear encoding and the latter relying on nonlinear recurrence, raises questions about the functional role of this distinction. In addition to representing spatial information, the hippocampus is essential for memory formation. Previous research has suggested a critical role for the recurrent connections in CA3 in storing and recalling associative memories (Rebola et al., 2017). We hypothesized that the distinct reliance on nonlinear recurrence by spatially and non-spatially tuned neurons could be an indicator of their involvement in perceptual versus mnemonic encoding.

To investigate this, we quantified the extent to which each CA3 neuron encodes information about past observations by assessing how well each neuron’s activity can be predicted by task-relevant features from prior time steps (Fig 5a). This analysis revealed distinct mnemonic profiles and firing patterns, with neurons differing in their selectivity for past versus present information (Fig 5c; Fig S10-S11). It identified that 48% of neurons were better predicted by stimuli currently perceived (classified as ’perceptual’, as this refers to information at present that is available to the senses), 22% were more selective to past information from recent time steps extending up to 10 steps into the past (near memory), and 29% encoded information from further in the past (distant memory, where memory refers to information from past events; Fig 5b). Moreover, neural predictivity dropped significantly when shuffling time steps within trial, suggesting specific temporal encoding rather than generic past encoding (Fig S12a, Wilcoxon signed-rank test: *P <* 10*^−^*^25^). Responses of memory neurons also showed little to no autocorrelation at their most predictive time steps, indicating that the most predictive time steps are unlikely to be due to stereotyped, event-locked activity patterns (Fig S12b).

**Figure 5:**
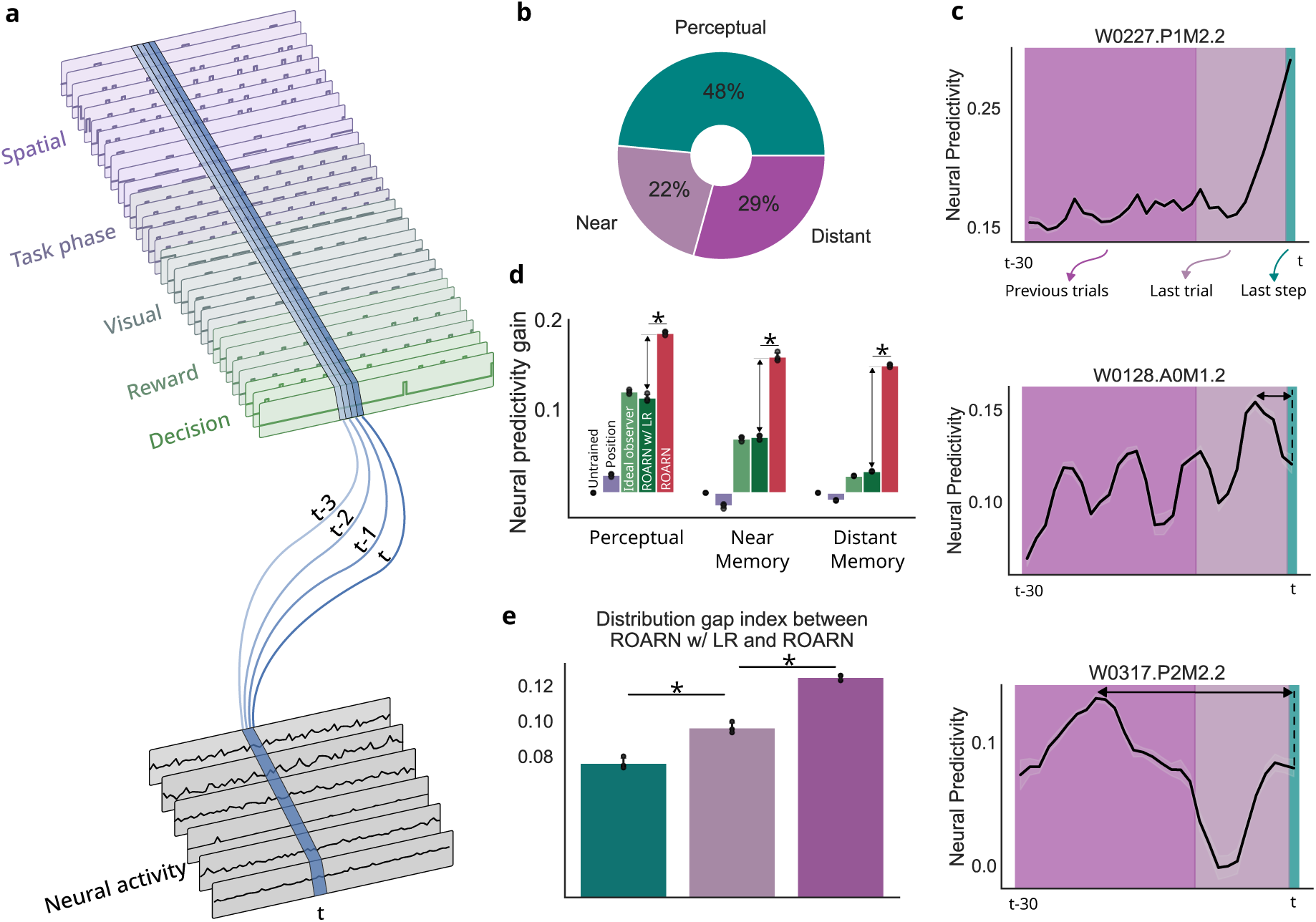
Mnemonic encoding. (a) Schematic of a linear mapping from task-relevant features to neuronal responses. A separate mapping is learned for each time step, allowing to investigate whether a neuron encodes past information. (b) Proportion of neurons classified per type of temporal encoding, indicating whether their activity is best predicted by past or present features. (c) Examples of mnemonic profiles for three neurons: Top – perceptual neuron, best predicted by current information; Middle – near-memory neuron, best predicted by features from recent steps; Bottom – distant-memory neuron, encoding information from previous trials. (d) Neural predictivity gain per temporal category, showing that ROARN outperforms ROARN w/ LR across all subsets of neurons (Wilcoxon rank-sum test: Perceptual *P <* 10*^−^*^8^, Near memory *P <* 10*^−^*^6^, Distant memory *P <* 10*^−^*^7^). (e) The gap in neural predictivity gain between ROARN and ROARN w/ LR increases for neurons encoding information from further in the past (Wilcoxon rank-sum test: Perceptual vs. Near memory *P* = 0.02, Near vs. Distant memory *P* = 0.02). All regressions were repeated across three random seeds. Mean *±* standard deviation over three random seeds

Reexamining the predictivity gap from prior analysis according to each neuron’s mnemonic classification revealed ROARN better predicts than ROARN w/ LR for every subset of neurons (Fig 5d Wilcoxon rank-sum test: Perceptual *P <* 10*^−^*^8^, Near memory *P <* 10*^−^*^6^, Distant memory *P <* 10*^−^*^7^; Fig S13). Moreover, as neurons encoded information from further in the past, the neural predictivity gap between both models became larger (Fig 5e Wilcoxon rank-sum test: Perceptual and Near memory *P* = 0.02 Near and Distant memory *P* = 0.02). ROARN w/ LR performed poorly in predicting memory neurons, whereas ROARN maintained better neural predictivity, resulting in a greater gap. This suggests that nonlinear recurrent connections play a crucial role in enabling memory neurons to integrate past information but provide more limited gains over linear combinations of task-related features for perceptual neurons. In other words, perceptual neurons are more readily explained by linear combinations of task-related features, while memory-related neural responses depend more on complex computational mechanisms, such as nonlinear recurrence.

### 2.5 Nonlinear mnemonic integration attenuates spatial tuning

In the rodent hippocampus, independent encoding schemes for spatial and episodic memory information have been observed, and it was suggested that nonlinearity may control the neurons sensitivity to which inputs are active (Leutgeb et al., 2005). Additionally, memory retrieval has been shown to modulate the spatial tuning of single neurons in the human entorhinal cortex, where some neurons exhibit robust memory-specific representations, while others display flexible spatial codes influenced by top-down memory demands (Qasim et al., 2019). Based on these findings, we hypothesized that the nonlinearity in memory neurons might disrupt spatial tuning. Our analysis revealed that neurons were less likely to exhibit spatial tuning as they encode information from further in the past. Specifically, 73% of perceptual neurons were spatially tuned, whereas only 21% of distant memory neurons had spatial tuning (Fig 6a), demonstrating a correlation between the temporal distance of encoded information and the strength of a neuron’s spatial selectivity (Fig 6b Pearson correlation with exponential fit: *r* = 0.51, S14). This discrepancy between space and time may stem from the role of memory neurons in integrating past information across multiple time steps and spatial positions. The differing predispositions to spatial tuning also account for the bimodal distribution of spatial response fields (Fig 2b), with one mode driven by perceptual neurons and the other by memory neurons (Fig 6c). Thus, processing information over extended periods may obscure spatial tuning, leading to two distinct neuron profiles: spatially-tuned neurons that linearly encode immediate perceptual information, and non-spatially tuned neurons that nonlinearly encode past information.

**Figure 6:**
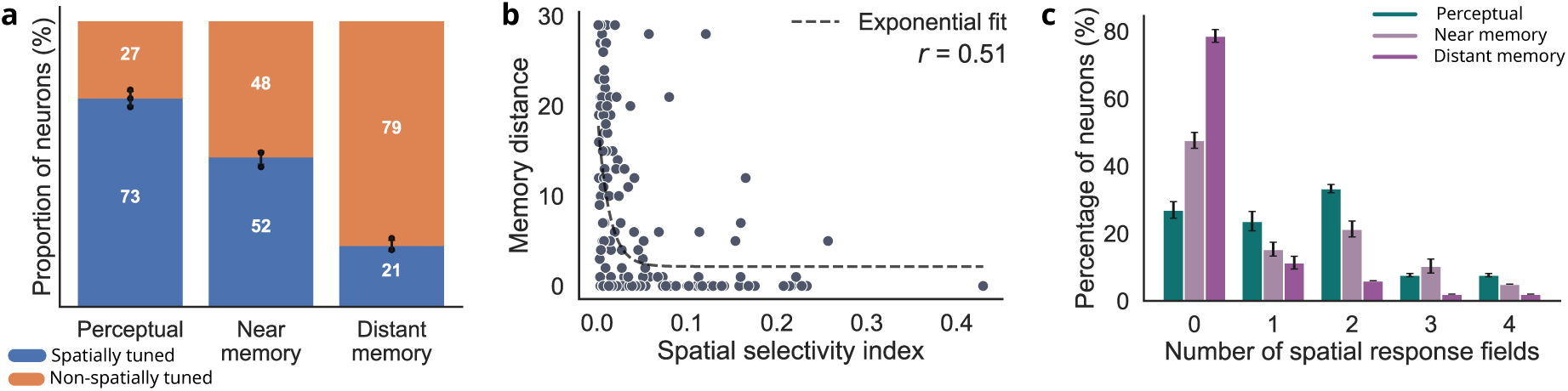
Relationship between spatial tuning and mnemonic encoding. (a) Proportion of neurons with spatial tuning across different temporal encoding categories, showing a decline in spatial tuning as neurons encode information further in the past. (b) Relationship between spatial selectivity and memory distance. An exponential fit to the data reveals a systematic decrease in spatial selectivity as neurons encode more distant past experiences (Pearson correlation with exponential fit: *r* = 0.51). (c) Distribution of spatial response fields across neuron subsets, indicating that memory neurons primarily contribute to the mode with no spatial response fields, while perceptual neurons drive the mode with two spatial response fields. All regressions were repeated across three random seeds. Bars represent mean *±* standard deviation over three random seeds

Overall, our findings reveal a continuum of hippocampal coding schemes that span from perceptual to memory-related representations, supported by a graded shift in computational complexity (Fig 7). At one end of this continuum, spatially tuned neurons exhibit linear response profiles that directly reflect current sensory input, making them well-suited for encoding perceptual states. At the other end, non-spatially tuned neurons rely on nonlinear computations to integrate information across extended time windows, enabling the encoding of more abstract, memory-related representations from the past. Intermediate neurons show a mixture of both properties, gradually transitioning from perceptual to memory-like responses. This gradient suggests that hippocampal representations are not confined to discrete cell types but instead emerge from a continuous spectrum of computations that flexibly support both present-moment information and temporally extended memory integration.

**Figure 7:**
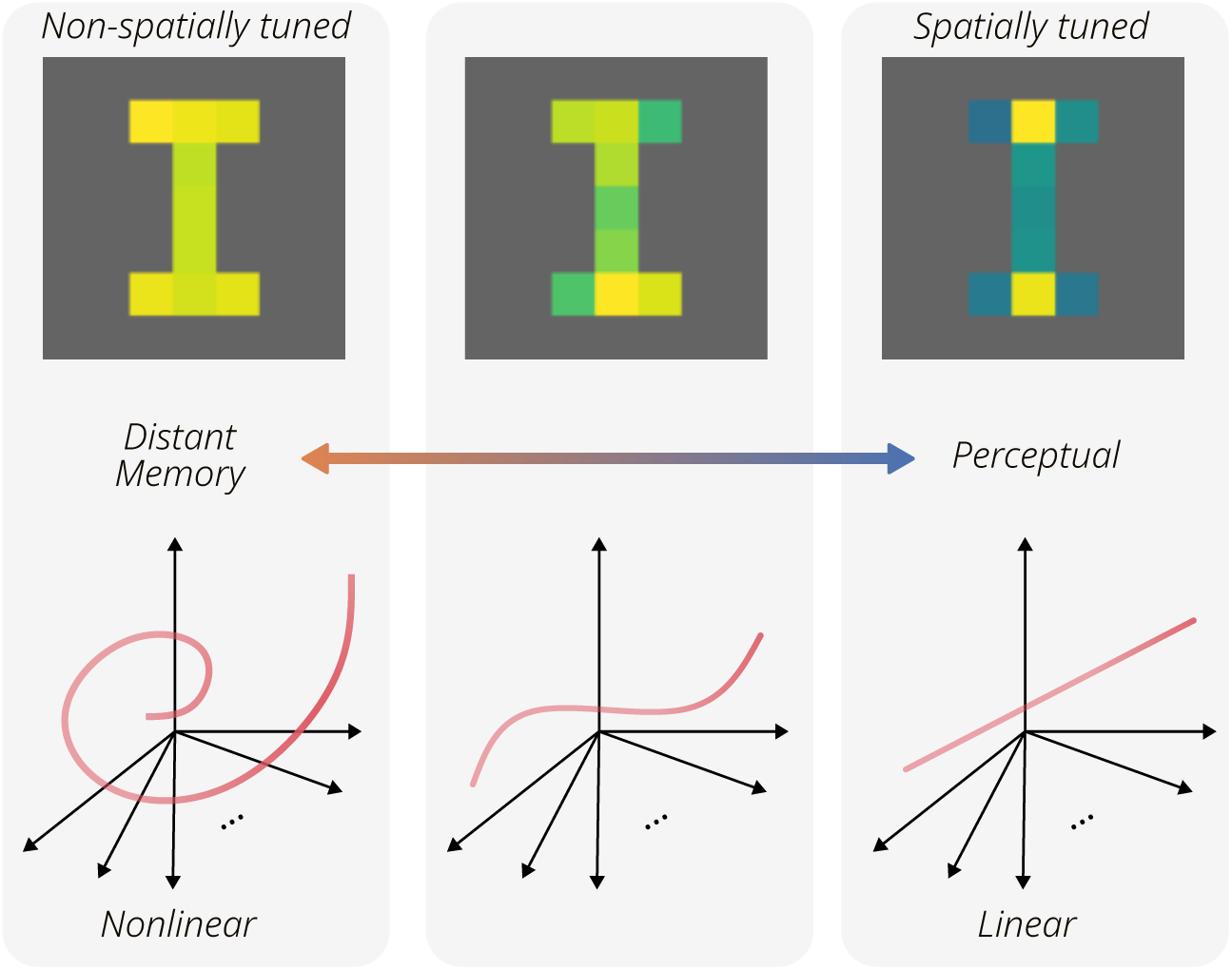
Relationship between spatial tuning, mnemonic encoding, and linearity or non-linearity of encodings. Hippocampal neurons represent both present and past information along a continuum of computational functions that vary in their degree of nonlinearity. Neurons that encode information about the present typically rely on more linear computations and are more likely to exhibit spatial tuning. In contrast, neurons that integrate information from the more distant past increasingly depend on nonlinear mechanisms, and as a result, are less likely to show spatial tuning.

## 3 Discussion

Using in-vivo electrophysiology and neural network modeling, we investigated the differences in information encoding between spatially and non-spatially tuned neurons in the primate hippocampus. Our findings suggest that spatial selectivity, or the lack thereof, can largely be attributed to whether neurons encode information in a linear or nonlinear manner. Furthermore, we observed that spatial selectivity decreases as neurons encode older information, indicating that neurons encoding present information are more likely to exhibit spatial tuning, while those involved in memory representation appear less spatially selective. Together, these observations highlight the diverse computational roles of CA3, illustrating how the region concurrently processes present inputs and integrates past experiences into memory via a spectrum of mechanisms spanning linear to nonlinear computation (Fig 7).

### Balancing Immediate Information and Memory Integration in CA3

CA3 neurons with linear response properties are well-suited to representing immediate information, as their activity can be rapidly decoded to guide real-time behavioral responses. Prior studies have shown that diverse stimuli, such as visual and somatosensory information, are encoded through linearly separable representations in both primate (Bernardi et al., 2020; Chang and Tsao, 2017) and rodent (Nogueira et al., 2023; Boyle et al., 2024) hippocampus. Linear perceptual encodings are thought to efficiently encode multiple variables, supporting compositional generalization to novel conditions (Courellis et al., 2024; Ito et al., 2022; Bakermans et al., 2025). Thus, linear encoding CA3 neurons create stable representations of immediate information, facilitating real-time behavioral adjustments to changing stimuli.

Conversely, CA3 neurons that encode past information nonlinearly appear to serve a distinct, complementary function, supporting complex, context-dependent processing essential for memory formation and recall. Nonlinear encoding also optimizes memory capacity by compressing high-dimensional experiences into efficient, selectively stored representations (Benna and Fusi, 2021; Spens and Burgess, 2024), enabling the hippocampus to retain a diverse range of information while maximizing storage efficiency. The functional diversity of neurons within CA3 appears to reflect a balance between immediate perceptual processing and the flexible retrieval of past experiences. By concurrently encoding linear and nonlinear representations, CA3 supports the integration of present information with long-term memory and facilitates the comparison between recent and temporally distant experiences (Howard and Eichenbaum, 2015). This dual representation allows CA3 to function as a dynamic processor, integrating diverse information across time and space to support adaptive behavior.

### Place Cells and Spatial Information

The classic view of hippocampal navigation posits that place cells primarily encode spatial information. However, recent studies challenge this perspective, revealing that hippocampal neurons often encode a mixture of spatial and non-spatial information. For instance, Stefanini et al. (2020) demonstrated that neurons traditionally considered non-spatial contribute equally to spatial encoding in CA1. Our study builds upon these findings and offers a unifying explanation for the paradoxical relationship between spatial selectivity and spatial information encoding. We show that macaque CA3 neurons encode a mixture of task-relevant variables, including position, along a continuum of linear and nonlinear functions. Neurons employing more nonlinear encoding mechanisms integrate information in ways that are generally less accessible by linear decoding approaches. Despite these differences, both spatially tuned and non-spatially tuned neurons contribute to spatial encoding, suggesting that neurons with sharp spatial tuning, such as place cells, are those utilizing linear mixing functions, while those with more uniform selectivity profiles employ nonlinear functions and are more likely to integrate past information.

### Task and Memory Dependence in Selectivity Profiles

Our findings establish a link between spatial selectivity and nonlinearity in neural encoding. However, an open question remains regarding when and why nonlinear versus linear functions are advantageous for encoding. The information content of hippocampal neurons have been shown to be dependent on task goals (Wikenheiser and Redish, 2015; Kelemen and Fenton, 2016; Gulli et al., 2020). Gulli et al. (2020) found that neuronal selectivity profiles changed drastically when macaque subjects explored the same virtual environment under different task instructions. Similarly, Markus et al. (1995) reported that place fields were more directional when rodents planned or followed a route between rewarded locations, as opposed to when rewards were randomly distributed, indicating that spatial selectivity in the hippocampus is influenced by behavioral demands. Finally, Eichenbaum et al. (1988); O’keefe and Conway (1980) showed that with simple variations to the task parameters (such as using two probes vs. one) one could make a task’s execution become hippocampal-dependent or independent. Our results predict that the distribution of spatial selectivity and, by extension, the degree of nonlinearity in encoding functions, mirrors behavioral requirements. Specifically, when animals engage in tasks that demand reliance on past information, the neural population is expected to exhibit a greater proportion of nonlinear, non-spatially tuned neurons. This suggests that hippocampal encoding mechanisms dynamically adjust based on cognitive demands.

### Cross-Region Differences

While our findings center on macaque CA3 neurons, it remains unclear whether similar encoding principles extend to other hippocampal subregions, such as CA1 or the subiculum. This raises important questions about how encoding properties are organized across the hippocampal circuit. One possibility is that connectivity between regions is structured by encoding type—for instance, CA3 neurons with linear, perceptual representations may preferentially project to similarly linear CA1 neurons. Under this scenario, the distribution of encoding types would be preserved downstream. Alternatively, if projections are pooled irrespective of encoding characteristics, downstream regions like CA1 may exhibit a greater proportion of neurons relying on nonlinear computations.

Recent work by Kong et al. (2024) identified a gradient within rodent CA3 itself: proximal CA3 (near the dentate gyrus) shows less recurrent dynamics and more consistent responses, while distal CA3 (closer to CA1) exhibits stronger recurrence and more context-dependent, variable responses. This gradient aligns with the continuum of linear perceptual to nonlinear mnemonic encoding observed in our study. Moreover, Barnes et al. (1990) reported that CA3 neurons are more spatially selective than their CA1 or subiculum counterparts, suggesting a stronger reliance on linear combinations of immediate features early in the hippocampal circuit. Meanwhile, studies like Lee et al. (2004) and Dong et al. (2021) have reported greater variability in CA1 responses, possibly reflecting increased nonlinearity and memory dependence in later processing. Together, these observations underscore the importance of examining how encoding properties are transformed—or preserved—across hippocampal subregions. Clarifying these cross-region differences will be key to understanding how the hippocampus supports both spatial and memory-related computations.

## Acknowledgments

We thank Drs. Mark Brandon, Adrien Peyrache, and Blake Richards for their helpful discussions and feedback on the manuscript. This research was supported by the Healthy-Brains-Healthy-Lives startup supplement grant, the NSERC Discovery grant RGPIN-2021-03035, and CIHR Project Grant PJT-191957. P.B. was supported by FRQ-S Research Scholars Junior 1 grant 310924, and the William Dawson Scholar award. M.D. was supported by the UNIQUE PhD Fellowship. All analyses were executed using resources provided by the Digital Research Alliance of Canada (Compute Canada) and funding from Canada Foundation for Innovation project number 42730.

## Supplementary Information

## A Methods

### A.1 Associative memory task

The task required monkeys to learn and apply context-dependent object-reward associations. Each trial began with the monkey at one end of the maze. During the context appearance task phase, one of two distinct textures (wood or steel) was applied to the maze walls, defining the current trial’s context. Upon reaching a central branched decision point in the maze, two of three uniquely colored disk-objects were presented, one in each of the two available arms. Monkeys had to learn a reward hierarchy associated with these objects, which was conditional upon the current wall texture (context). For example, in the ’wood’ context, object A might be high-reward, B medium-reward, and C low-reward, while in the ’steel’ context, this hierarchy is reversed (e.g., C *>* B *>* A).

Crucially, a new set of three colored disks was used for the objects in each recording session, and the assignment of these colors to the high, medium, or low reward positions within each context was randomized. This required the monkeys to learn new conditional associations between the context and object reward values daily. Monkeys made a choice by navigating towards one of the displayed objects to receive a juice reward commensurate with the object’s value in the current context. Each trial was divided into five task phases: trial start, precontext, context appearance, object appearance, and object approach (for more details, see Gulli et al. (2020)).

### A.2 ROARN

The ROARN model is optimized using the Proximal Policy Optimization (PPO) reinforcement learning algorithm (Schulman et al., 2017). The model’s architecture includes a Convolutional Neural Network (CNN) for processing visual input (LeCun et al., 1989), a Long Short-Term Memory (LSTM) (Hochreiter and Schmidhuber, 1997) network for capturing temporal dependencies, and a policy network for decision-making. The model receives 5x5 visual inputs from the Minigrid environment (Chevalier-Boisvert et al., 2023). To facilitate model convergence, the number of trials per episode is gradually increased, starting from 10 trials and scaling up to 350 trials. The action space consists of three possible actions: turning left, turning right, and moving forward. In the macaque monkey experiments, the association between context cues and goal objects changes between each recording session. To indicate these session changes to the agent, a session indicator is provided as a one-hot vector.

During the training phase, the ANN model parameters are optimized to maximize the expected reward across trials. Through this process, the convolutional network which loosely mimics the animal’s visual system learns object level representations from the learned sensory inputs, while the recurrent neural network learns to integrate information across multiple time steps and to associate context cues with goal objects that lead to larger rewards more frequently. After training, the Reward Optimized Artificial Recurrent Network (ROARN) learns to perform the optimal behavior in most trials (Fig S4).

### A.3 Neural predictivity

To predict hippocampal neural activity, we employed a previously used approach for comparing ANN activations to the neural activity of the visual (Yamins et al., 2014), motor (Sussillo et al., 2015), and auditory cortices (Kell et al., 2018). We subjected the artificial agent to the same trial sequences experienced by the animal subjects. We then performed a linear regression analysis using a linear SVM (Support Vector Machine) (Cortes and Vapnik, 1995) to predict the neurons’ firing rates from the model’s unit activations. We used 10-fold cross-validation where the time series is divided in 10 temporally contiguous periods of equal size in terms of number of datapoints, and we trained the SVM using the data from 9 of them and tested on the remaining data (Stefanini et al., 2020). The neural predictivity score was obtained by computing the Pearson correlation, *ρ*, between predicted and actual firing rates. The neural predictivity gain of a model is the neural predictivity adjusted by the untrained ROARN score to report the actual neural predictivity gained upon a random baseline obtained from processing the same inputs with random weights.

### A.4 Spatial response fields

To determine whether a neuron or unit exhibited selective firing in any of the nine distinct maze areas (four arms, two branches, and three corridor sections), we followed the protocol from Gulli et al. (2020) and employed a statistical permutation test based on spatial position and firing rates. For each trial, a vector of spatial positions was created, and the neuron’s firing rate was computed at each position. These vectors were aggregated across trials to generate an average firing rate for each spatial bin. To assess whether the observed firing rates were significantly elevated compared to chance, we performed a circular shuffling procedure. This shuffling process was repeated 1,000 times to generate a null distribution of firing rates for each bin. A bin was considered statistically significant if its empirical firing rate exceeded the 99th percentile of the null distribution (p-value = 0.01), with false discovery rate (FDR) correction applied using the Benjamini–Hochberg procedure. Spatial response fields were defined as any of the nine maze areas containing at least one statistically significant bin. Neurons or units with at least one spatial response field were classified as spatially tuned.

### A.5 Encoding of spatial and non-spatial features analysis

The ideal observer model incorporates all relevant features, including spatial position, direction of travel, visual cues, selected goal-object, reward value received, and task phases. A linear Support Vector Machine (SVM) is used to predict the firing rates of a neuron based on the ideal observer representation. We assess the contribution of each feature by examining the absolute and normalized weights of the fitted SVM on the individual features encoded in the ideal observer. Spatial features consist of the animal’s spatial position and direction of travel, while task phases encompass the five trial stages: Trial Start, Precontext, Context Appearance, Object Appearance, and Object Approach (decision). Non-spatial features include all other features. A feature is deemed important for predicting a neuron’s responses if its weight is at least two standard deviations above the average weight.

### A.6 Mnemonic properties

Mnemonic properties of a neuron are quantified by assessing how well features from previous time steps predict its current activity. Neural predictivity is computed similarly to the ideal observer model, except that the model is trained using information from earlier time steps. To determine the most predictive time steps for each temporal category, each neuron’s temporal profile is smoothed using a moving average window of size 3 and then averaged across random seeds. The time step with the highest neural predictivity within the near or distant memory range is then compared to the perceptual neural predictivity using a Wilcoxon rank-sum test. Memory distance is defined as the temporal gap between the most predictive time step and the present moment. A memory distance of 0 indicates that the neuron is best predicted by currently perceived features, classifying it as a perceptual neuron. Greater memory distances indicate a preference for older features, classifying the neuron as a memory neuron. Neurons encoding past information from up to one trial ago (within the past ten time steps needed to traverse the maze) are categorized as near memory neurons. Neurons encoding information from further in the past (up to 30 time steps) are categorized as distant memory neurons. Memory strength is defined as the difference in neural predictivity between the most predictive time step and the perceptual time step.

### A.7 Spatial selectivity index

We assessed spatial variance to determine if the variance in firing rates is primarily due to spatial selectivity. Spatial variance is calculated as the ratio of the variance of the mean firing rates at each spatial position to the overall variance of all firing rates.

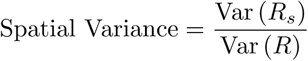

where:

- *R_s_* is the set of mean firing rates at each spatial position, defined as *R_s_* = *{µ*_1_*, µ*_2_*, . . ., µ_N_ }*,

where: 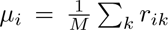 with *r_ik_* representing the firing rate observed at position *i* across *M* trials.

- *R* is the set of all firing rates across spatial positions and trials, i.e., *R* = *{r_ik_}*.

Therefore, Var(*R_s_*) is the variance of the mean firing rates across spatial positions and Var(*R*) is the overall variance of all firing rates *r_ik_* across all spatial positions and trials. This ratio captures the variance between spatial positions normalized by the overall firing rate variance. If a neuron fires selectively based on spatial position, the firing rates will differ significantly between spatial positions, resulting in higher variance between spatial positions. Consequently, a larger portion of firing rate variance will be attributed to spatial selectivity, leading to a higher spatial variance which indicates that spatial position strongly influences the firing rate. Notably, this metric correlated with the number of spatial response fields (Fig S15)

## B Supplementary figures

**Figure S1:**
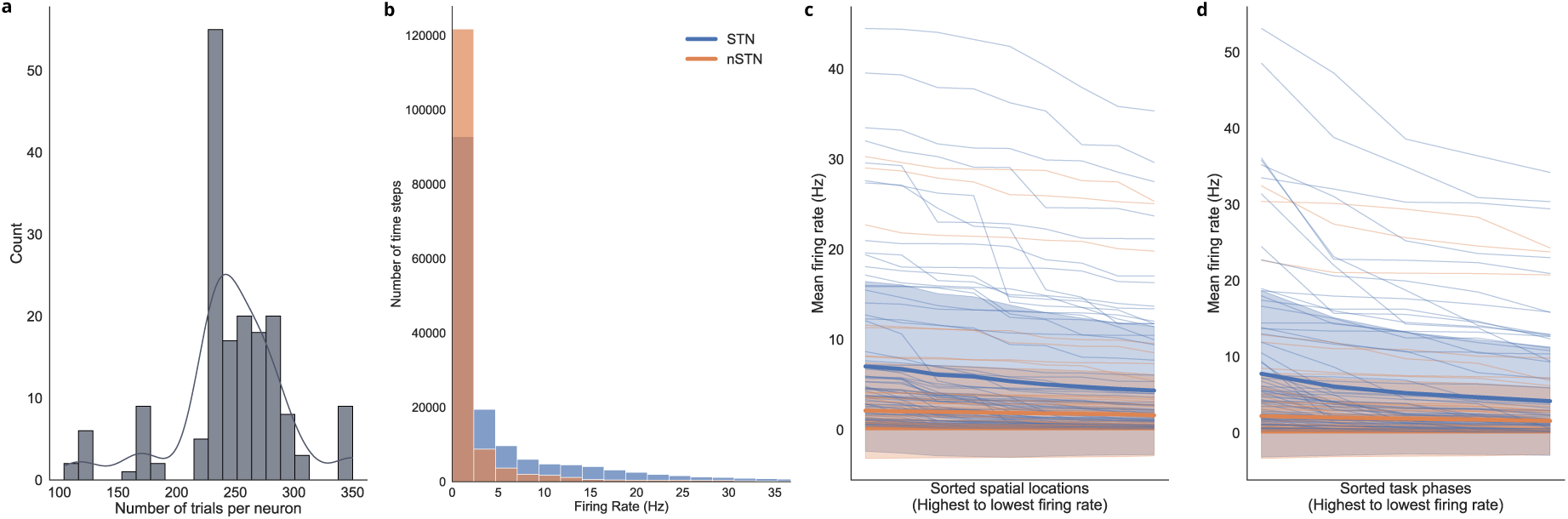
Neural properties. (a) Distribution of the number of trials recorded per neuron. Each bar represents the count of neurons with a given number of recorded trials, with an overlaid kernel density estimate illustrating the overall distribution. (b) Distribution of firing rates across time steps. To match the resolution of the computational models, firing rates are computed at each discrete time step corresponding to the model’s environment. Spatially tuned neurons (STNs) are shown in blue, while non-spatially tuned neurons (nSTNs) are in orange. Outlier values are excluded for clarity. (c) Mean firing rates of neurons across different spatial locations, sorted from highest to lowest firing rate for each neuron. Bold lines represent the mean firing rate across all STNs and nSTNs, respectively. (d) Mean firing rates of neurons across different task phases, sorted in descending order of firing rate for each neuron. Similar to (c), STNs and nSTNs are distinguished by color, with bold lines indicating the mean firing rates within each group.

**Figure S2:**
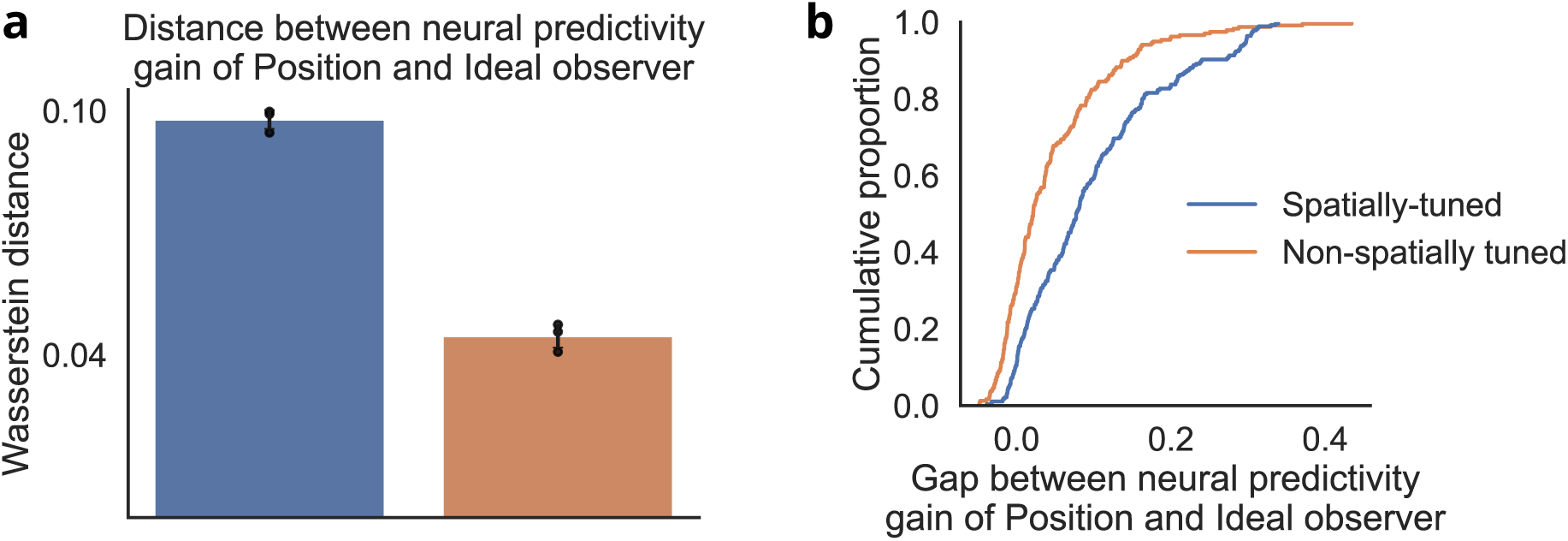
Prediction of spatially tuned neurons and non-spatially tuned neurons responses with only spatial features or all task-relevant features. (a) Wasserstein distance between neural predictivity gain of Position and Ideal observer. Predictions from Ideal observer are significantly different than those from solely spatial position features for STNs, but the additional features resulted in a smaller improvement for nSTNs. (b) Empirical cumulative distribution of the gap between the neural predictivity gain of Position and Ideal observer for each neurons. The gap between both model was smaller for nSTNs compared to STNs, two-sample Kolmogorov–Smirnov test *P <* 10*^−^*^12^) Mean *±* standard deviation over three random seeds

**Figure S3:**
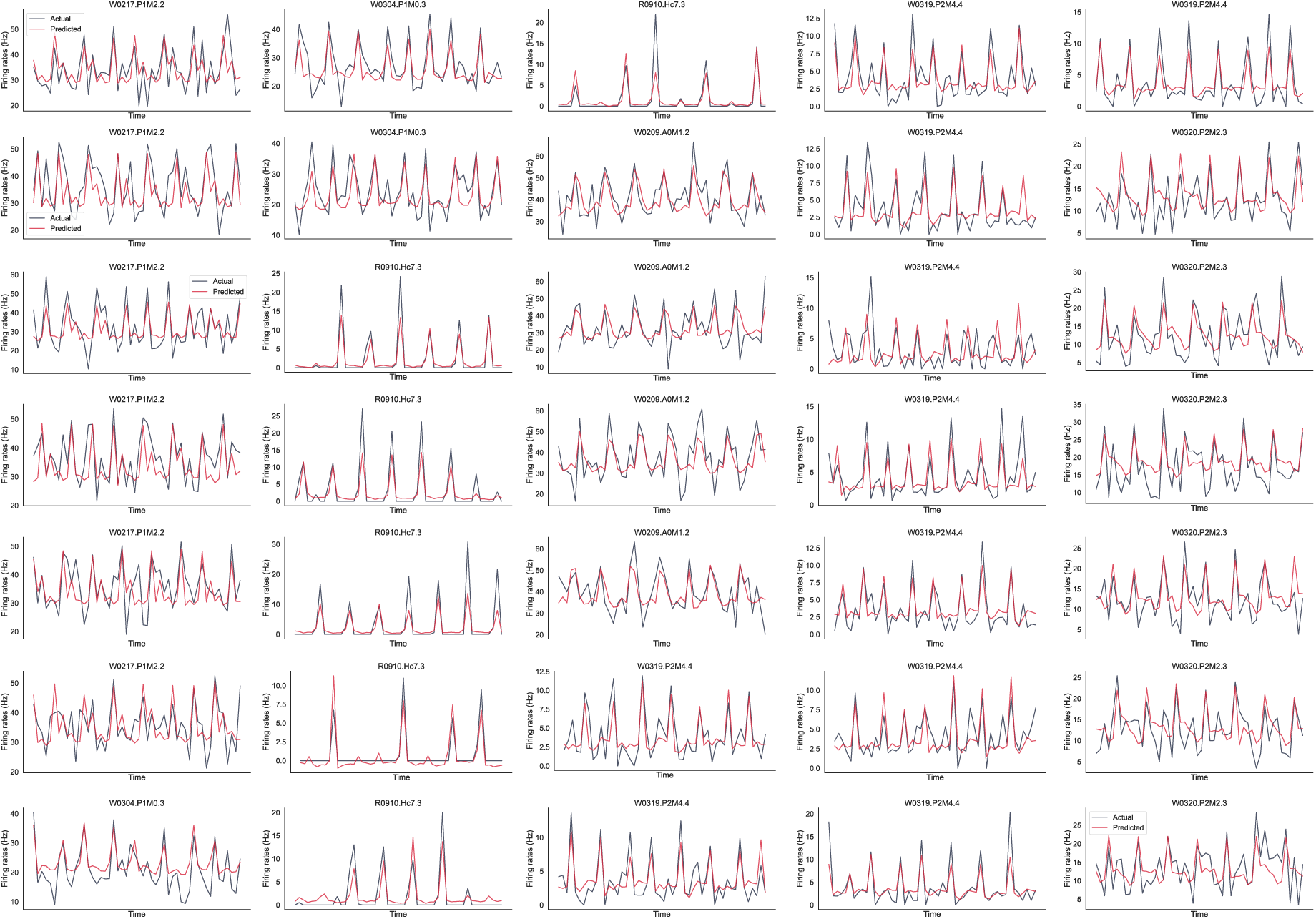
Firing rate predictions. Example of hippocampal neuron responses (black) and their corresponding predictions from the ROARN model (red). Examples are selected from six well-predicted neurons and are of the duration of 50 time steps.

**Figure S4:**
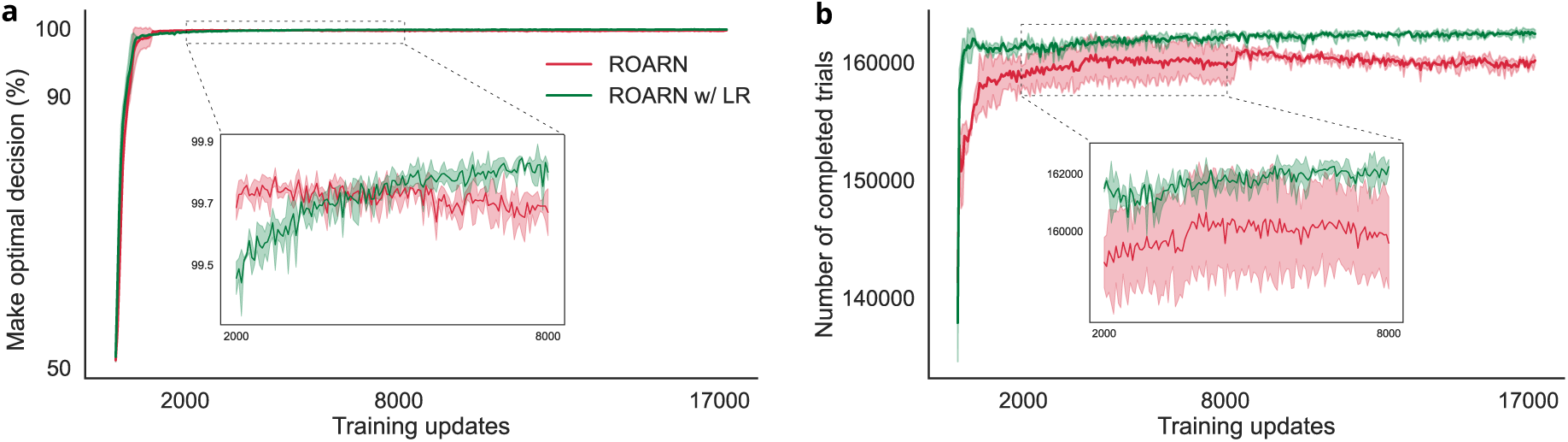
Learning curves. (a) ROARN and ROARN w/ LR achieve similar levels of optimality in navigating to the best goal object available throughout training. Both ROARN and ROARN w/ LR rapidly improve their decision-making performance early in training, reaching near-optimal behavior. Throughout training, both models maintain similar levels of optimality, making the correct decision with high accuracy. The inset shows minor difference between models. (b) The number of completed trials per fixed amount of time steps follows a similar trajectory for both models, indicating that both models achieve comparable navigation proficiency. Mean *±* standard deviation over three random seeds.

**Figure S5:**
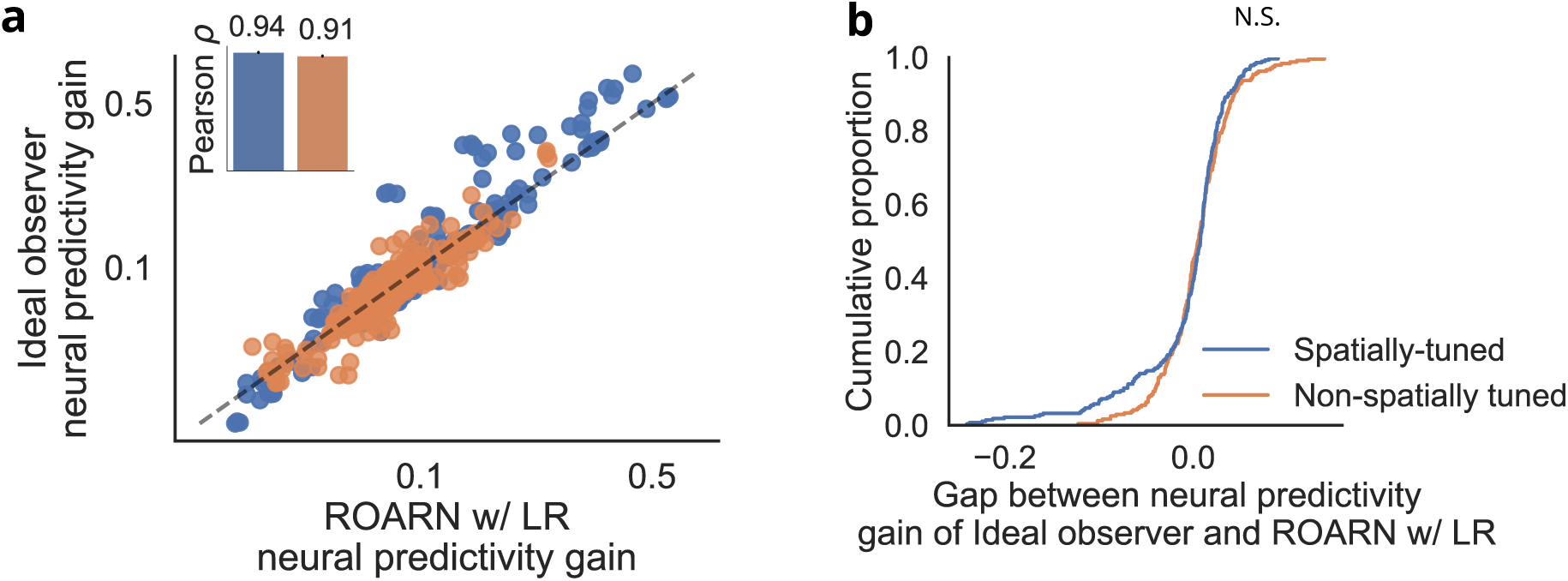
Reliance on nonlinear recurrence to capture nSTNs. (a) Scatter plot comparing neural predictivity gain between ROARN w/ LR and the ideal observer. Removing the nonlinear recurrence makes prediction of neural responses similar to those of the ideal observer. (b) Empirical cumulative distribution function of the gap between the neural predictivity gain of ROARN w/ LR and ideal observer for each neurons (two-sample Kolmogorov–Smirnov two-sided test *P* = 0.20).

**Figure S6:**
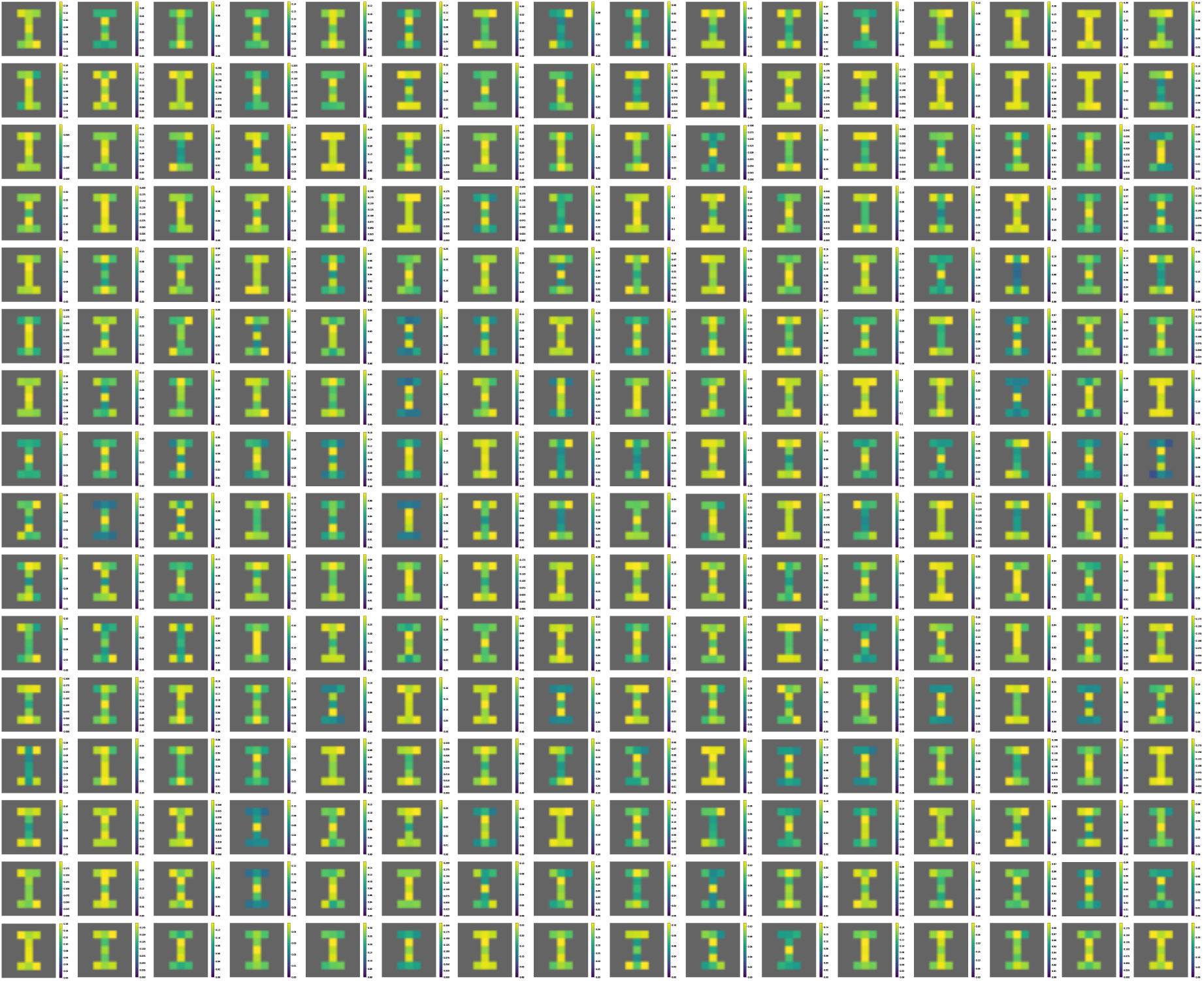
Untrained ROARN spatial maps. Spatial maps of the 256 units in the recurrent layer of an untrained ROARN with random weights.

**Figure S7:**
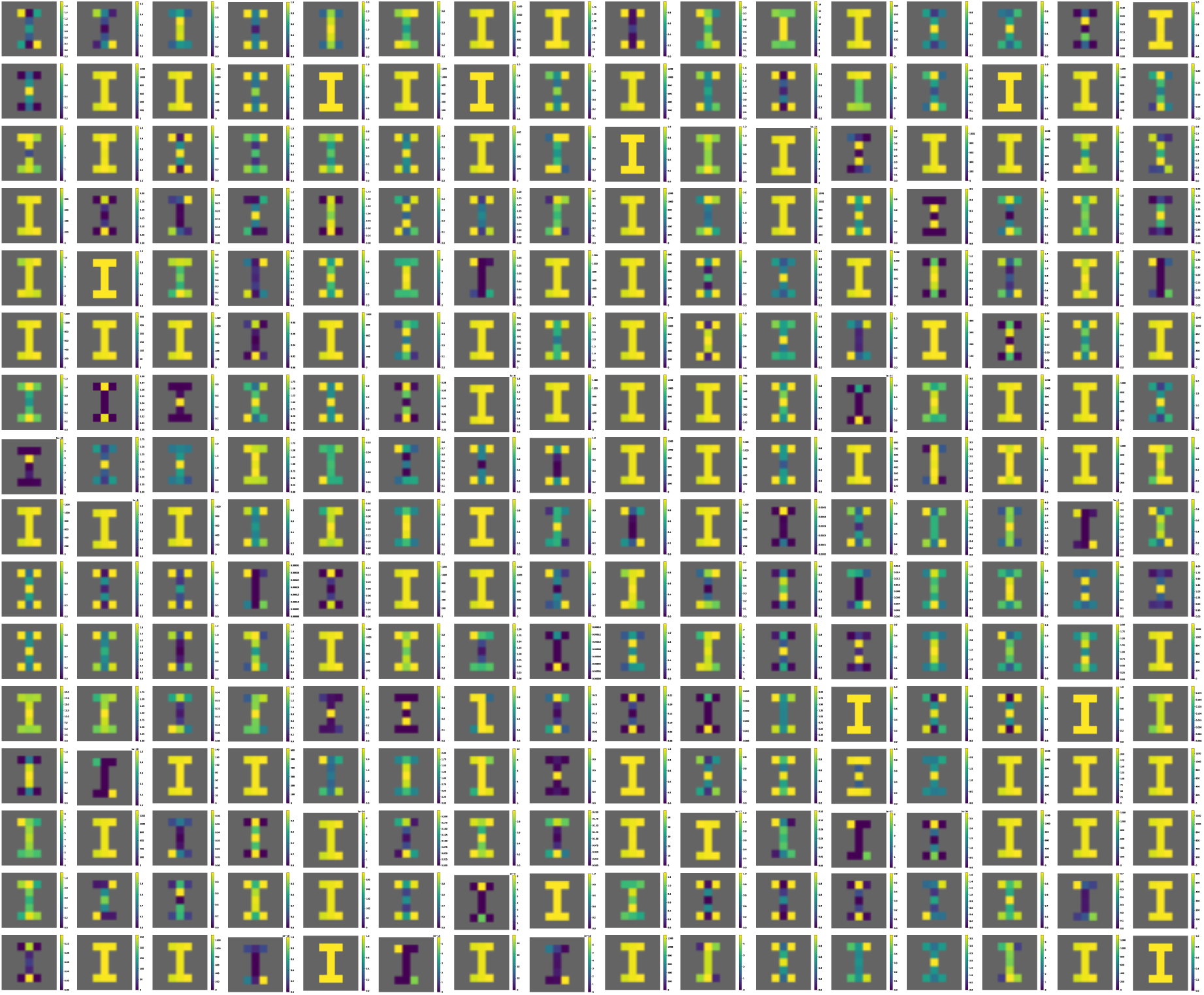
ROARN spatial maps. Spatial maps of the 256 units in the recurrent layer of ROARN.

**Figure S8:**
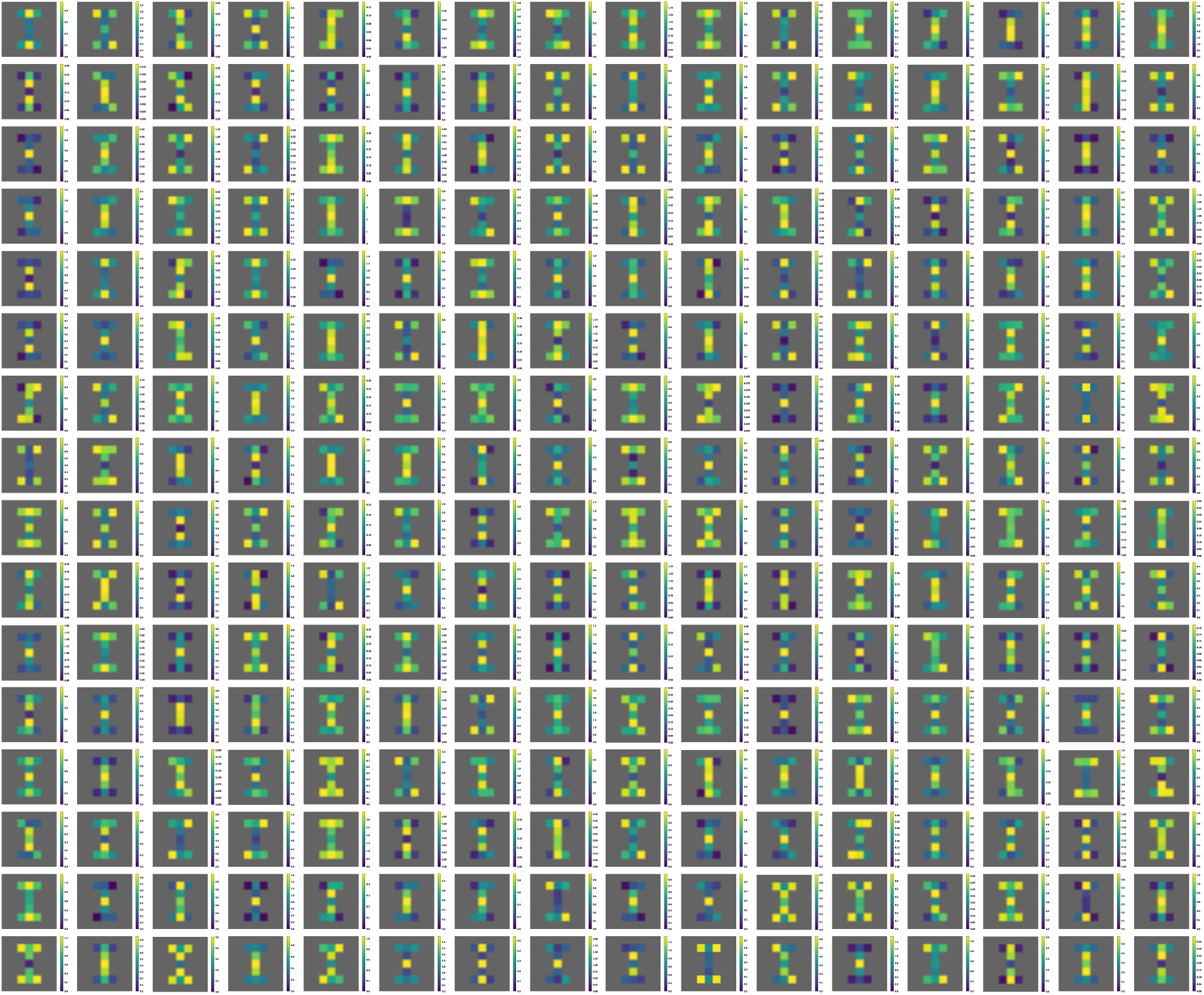
ROARN w/ LR spatial maps. Spatial maps of the 256 units in the linear recurrent layer of ROARN w/ LR.

**Figure S9:**
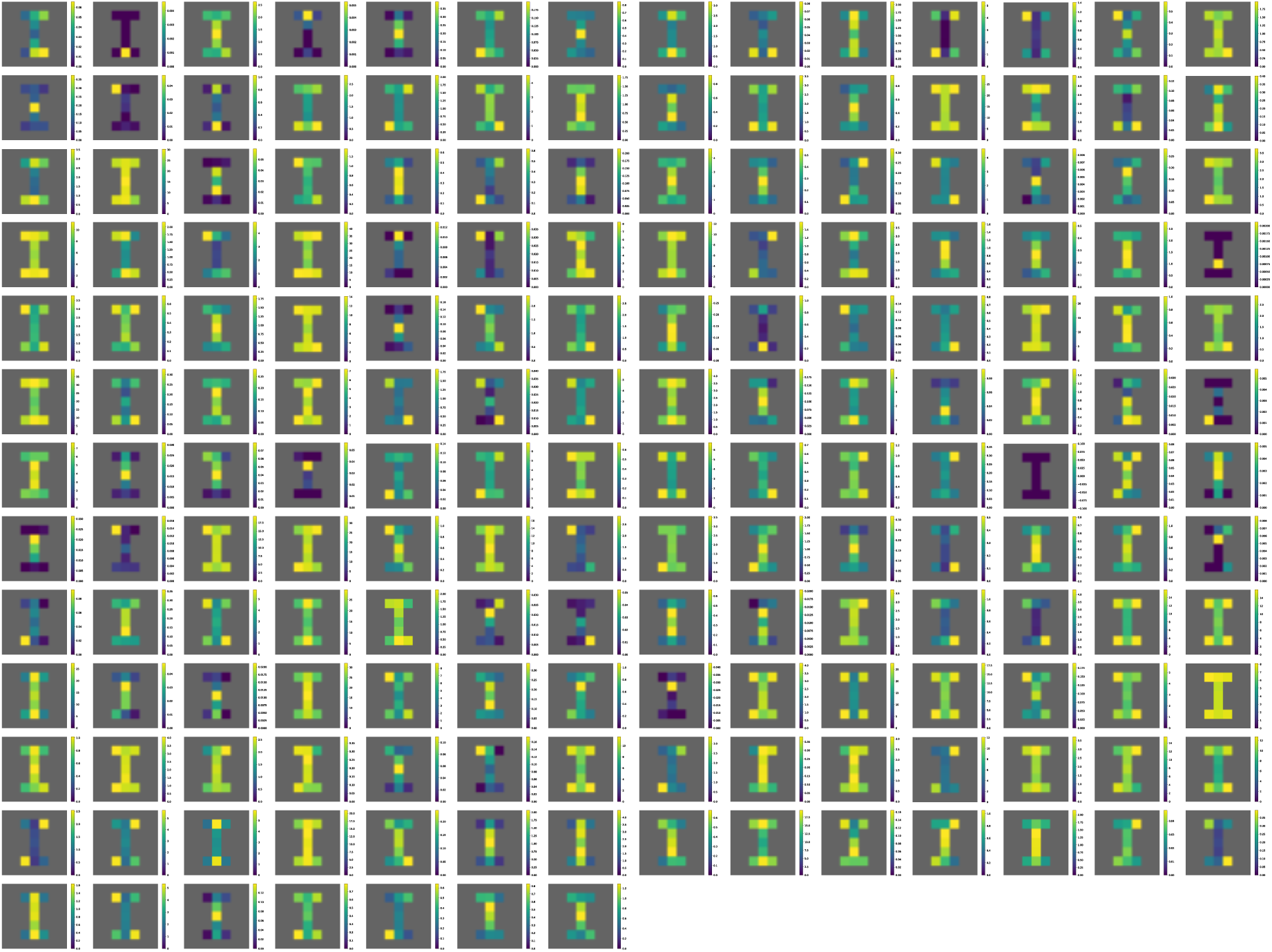
Spatial maps. Spatial maps of the 175 hippocampal neurons.

**Figure S10:**
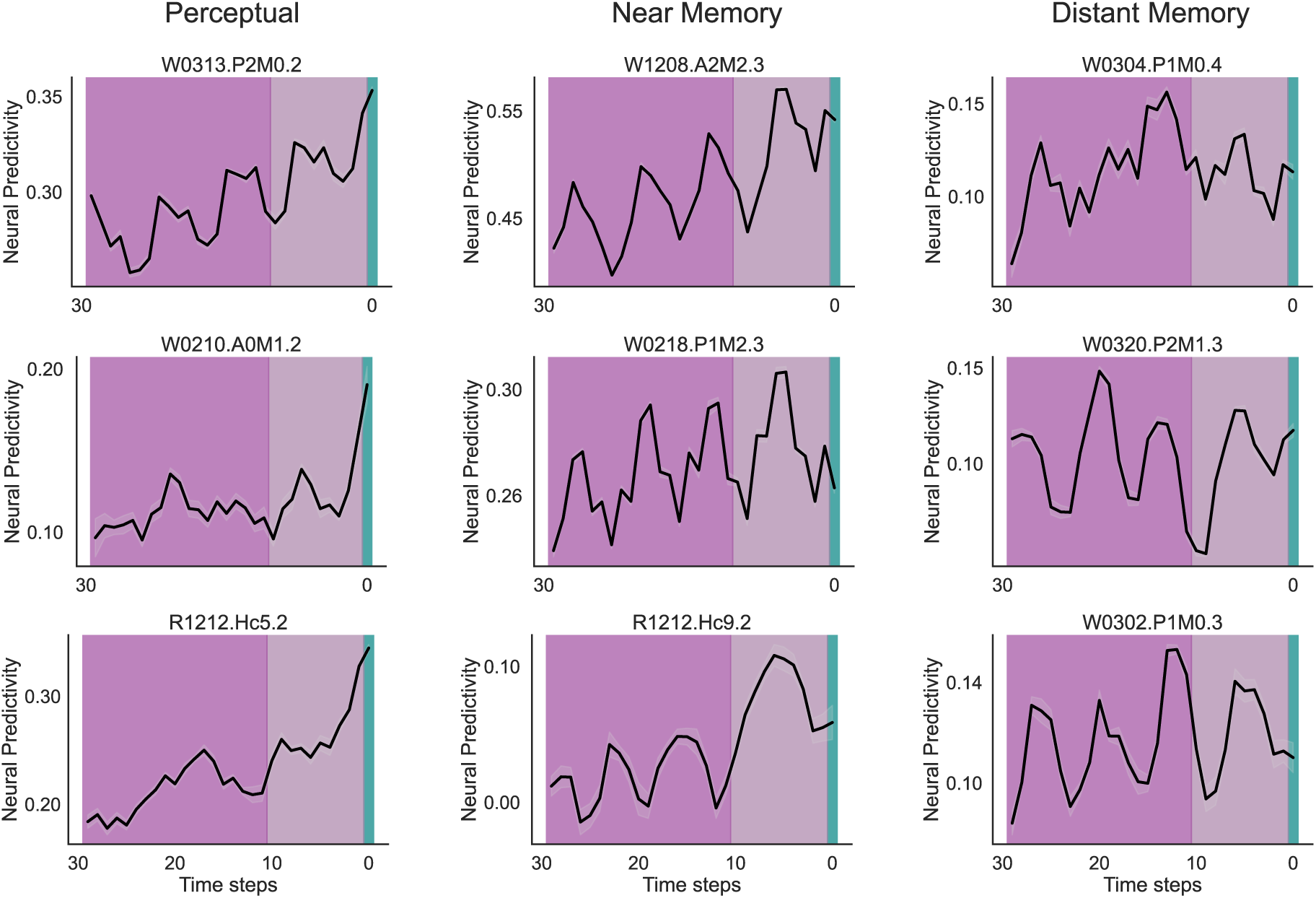
Mnemonic profile examples. Examples of mnemonic profile of perceptual, near memory, and distant memory neurons (from left to right)

**Figure S11:**
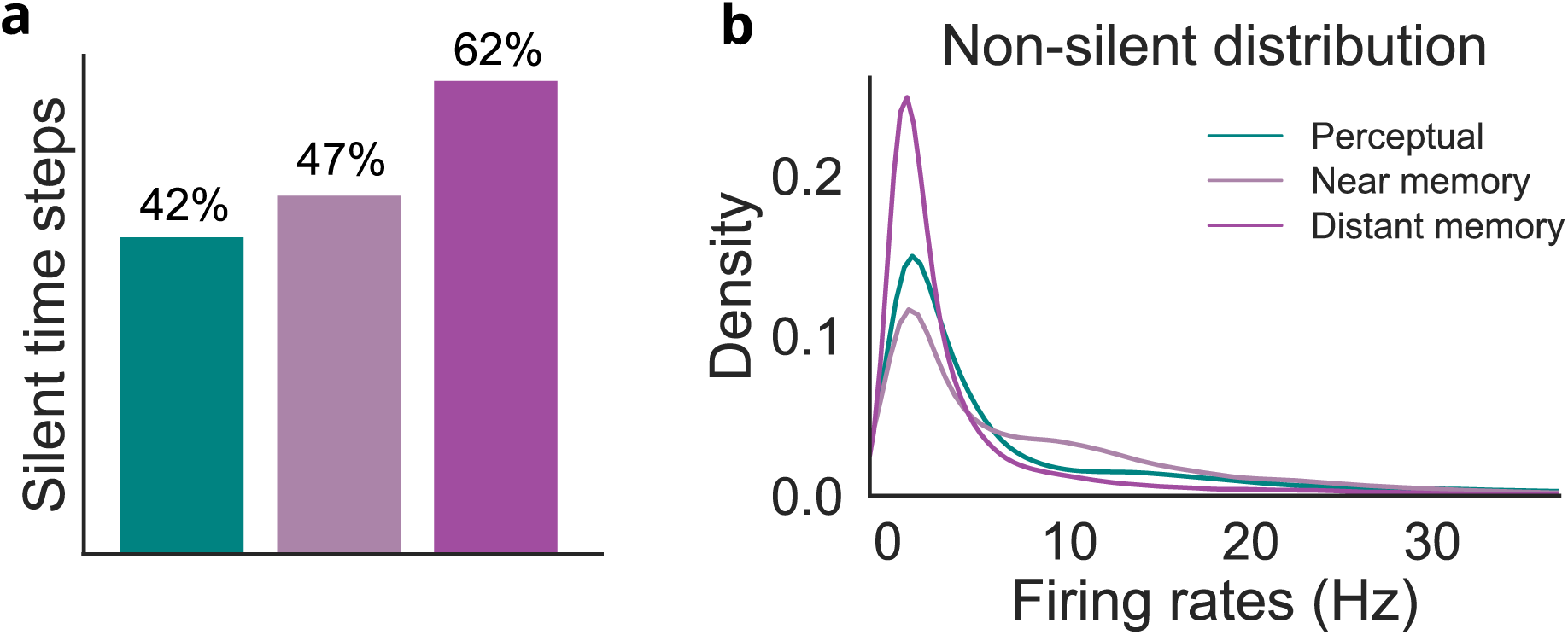
Firing rate and temporal encoding. (a) Proportion of time steps in which neurons remain silent (i.e., exhibit no spiking activity), showing that distant memory neurons fire less frequently than perceptual and near memory neurons. (b) Probability density distribution of firing rates during active time steps (non-silent periods), separately shown for perceptual, near memory, and distant memory neurons. The distributions highlight differences in firing dynamics, with distant memory neurons exhibiting lower mean firing rates compared to the other categories.

**Figure S12:**
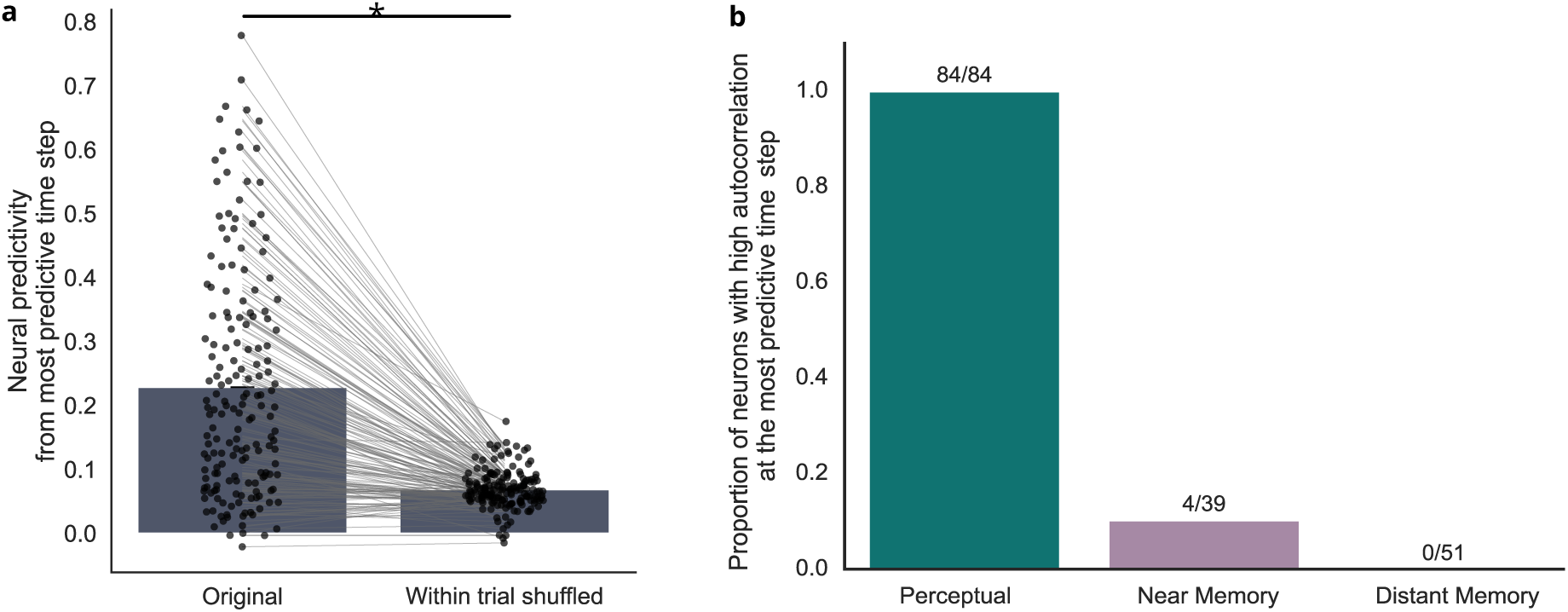
Mnemonic encoding controls. (a) Neural predictivity from the most informative past time step significantly decreases when that time step is replaced with a randomly selected time step from the same trial (Wilcoxon signed-rank test: *P <* 10*^−^*^25^), indicating temporal specificity. (b) Proportion of neurons within each category for which the most predictive time step shows significantly higher autocorrelation with the current neural activity (t) than other past time steps (t–m), based on a Wilcoxon rank-sum test (threshold *P <* 0.05). While all perceptual neurons (84/84) showed high autocorrelation, only 4/39 near memory and 0/51 distant memory neurons had significant autocorrelation, suggesting that the most predictive time steps are not due to stereotyped, event-locked firing patterns.

**Figure S13:**
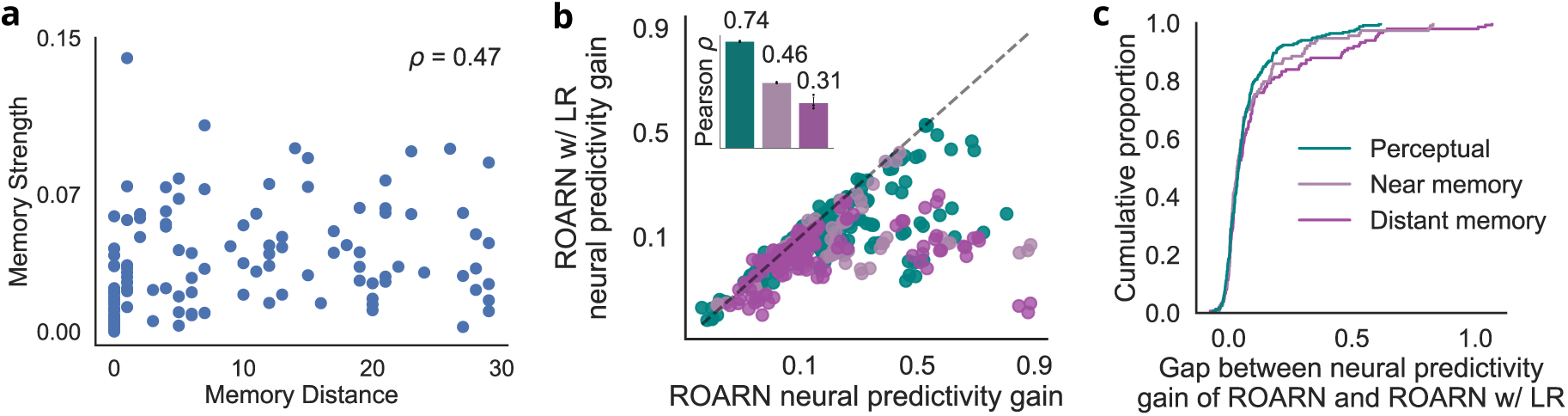
Mnemonic propesties. (a) Relationship between memory distance and memory strength, where memory strength is the difference in neural predictivity between the current time steps and the maximum neural predicitivty across all time steps. (b) Scatter plot comparing neural predictivity gain between ROARN w/ LR and ROARN. (c) Empirical cumulative distribution of the gap between the neural predictivity gain of ROARN with and without nonlinear recurrence.

**Figure S14:**
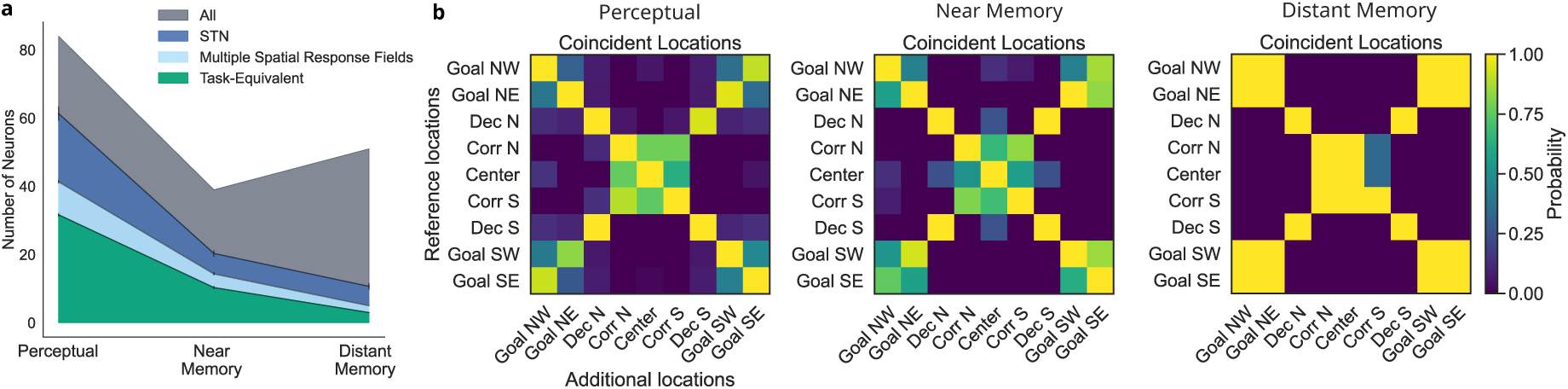
Spatial tuning properties and temporal encoding. (a) Number of neurons with at least one spatial response field, multiple spatial response fields, and their spatial response fields distributed at task-equivalent locations (b) Coincident locations of spatial response fields for perceptual, near memory, and distant memory neurons with multiple response fields, demonstrating symmetrical tuning properties. Mean ±standard deviation over three random seeds

**Figure S15:**
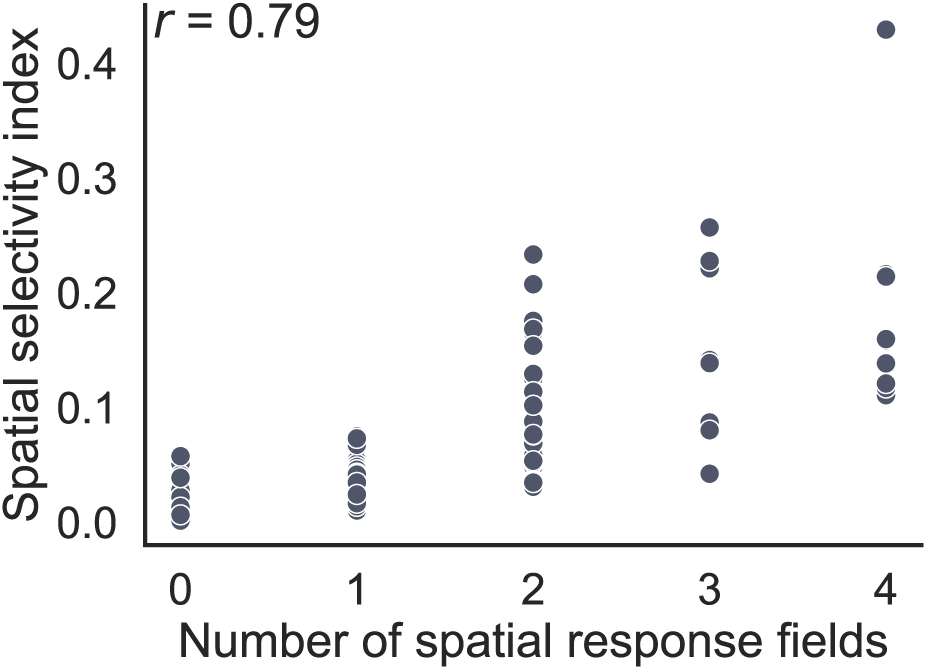
Relationship between number of spatial response fields and spatial selectivity index.

## C Image attribution

RNN schema in Figure 3a is modified from scidraw.io/drawing/495, shared under creative commons license (CC-BY) by Federico Claudi. Monkey schema in Figure 2a and 3a is modified from scidraw.io/drawing/445, shared under creative commons license (CC-BY) by Andrea Colins Rodriguez. Computer screen icon in Figure 2a and 3a is modified from thenounproject.com/icon/computer-7392374/, shared under creative commons license (CC-BY) by Jae Deasigner. Joystick icon in Figure 2a and 3a is modified from thenounproject.com/icon/joystick-7337812/, shared under creative commons license (CC-BY) by Rikas Dzihab.

